# Light-induced nanoscale deformation in azobenzene thin film triggers rapid intracellular Ca^2+^ increase via mechanosensitive cation channels

**DOI:** 10.1101/2022.09.27.509666

**Authors:** Heidi Peussa, Chiara Fedele, Huy Tran, Julia Fadjukov, Elina Mäntylä, Arri Priimägi, Soile Nymark, Teemu O. Ihalainen

## Abstract

Epithelial cells are in continuous dynamic biochemical and physical interaction with their extracellular environment. Ultimately, this interplay guides fundamental physiological processes. In these interactions, cells generate fast local and global transients of Ca^2+^ ions, which act as key intracellular messengers. However, the mechanical triggers initiating these responses have remained unclear. Light-responsive materials offer intriguing possibilities to dynamically modify the physical niche of the cells. Here, we use a light-sensitive azobenzene-based glassy material that can be micropatterned with visible light to undergo spatiotemporally controlled deformations. The material allows mechanical stimulation of single cells or multicellular assemblies, offering unique opportunities for experimental mechanobiology. Real-time monitoring of consequential rapid intracellular Ca^2+^ signals reveal that Piezo1 is the key mechanosensitive ion channel generating the Ca^2+^ transients after nanoscale mechanical deformation of the cell culture substrate. Furthermore, our studies indicate that Piezo1 preferably responds to lateral material movement at cell-material interphase rather than to absolute topographical change of the substrate. Finally, experimentally verified computational modeling of the signaling kinetics suggests that the lateral mechanical stimulus triggers multiplexed intercellular signaling that involves Na^+^, highlighting the complexity of mechanical signaling in multicellular systems. These results give mechanistic understanding on how cells respond to material dynamics and deformations.

## Introduction

Cells are constantly subjected to mechanical stimuli such as stretch, compression, osmotic stress, and shear (Yang et al. 2017; Sapir and Tzlil 2017). The mechanical information cells receive from their physical environment co-regulates their form and functions, thus allowing cells to adapt to their niche. (Wang and Li 2010; Iskratsch, Wolfenson, and Sheetz 2014; Martino et al. 2018) The physical cues are often sensed by specific and highly dynamic protein complexes at the cell membrane, e.g., integrin-rich focal adhesions (cell-extracellular matrix (ECM) junctions), cadherin-based adherens junctions (cell-cell junctions), and mechanosensitive ion channels on the cell membrane (Martino et al. 2018; Eyckmans et al. 2011; Jin, Jan, and Jan 2020; Douguet and Honoré 2019; Kefauver, Ward, and Patapoutian 2020). These mechanosensing structures form an important signal transducing interface between cells and their physical environment influencing physiological processes. This interface allows cells to sense e.g., ECM rigidity and topography, and spatial changes in these characteristics can trigger directed cellular migration along rigidity gradient (durotaxis) (Lo et al. 2000) or along topography gradient (topotaxis) (Park, Kim, and Levchenko 2018). Also temporal changes in this interface affect cell behavior and therefore, the dynamics of the cell-material interaction is of immense importance (Hoffman, Grashoff, and Schwartz 2011). Defects and misregulation at the cell-material interface can also play a role in pathological conditions. For example, changes in the microenvironment promote epithelial-mesenchymal transition, a process associated to cancer metastasis (Scott, Weinberg, and Lemmon 2019; Davis et al. 2014). Therefore, the physical characteristics of cell-material interface and its effects on cellular physiology are highly important.

The physical properties of the cell-ECM interface can be actively manipulated by using different materials (Y. Li, Xiao, and Liu 2017) and over the last 20 years the number of techniques to characterize cellular responses to the applied mechanical stimulation has exploded (Xi et al. 2019). However, the existing solutions mostly apply basal mechanical stimulation to the entire population, whereas options to reproducibly induce mechanotransductive events at single cell or subcellular levels remain limited. In this context, the manipulation of material surface topography is a highly promising approach to study the interface between cells and the ECM (Ventre, Causa, and Netti 2012). Different micro-and nanoengineered topographies can stimulate collective migration of cells, guide axonal growth of neurons promoting their differentiation, and influence migration of cancer cells (Song et al. 2016; Londono et al. 2014). These studies have mostly been conducted with static topographies that cannot be modified after fabrication. However, the ECM is constantly being assembled, disassembled, and reorganized in micro- and nano-scale (Larsen et al. 2006; Lu et al. 2011) to adapt to the new environmental conditions. Biomimicking the dynamic nanoscale behavior of the cell-ECM interface has proven to be difficult and current understanding of what kind of topographical cues or forces cells can sense and how it is mechanistically achieved is limited. For this, azobenzene-based, light-controllable materials offer a powerful approach. Azobenzene-containing materials have been recently proposed as smart biointerfaces, as their topography can be precisely manipulated via visible light. Azobenzenes can isomerize in response to light between *trans* and *cis* forms, characterized by distinct molecular geometries. This switching process endows glassy materials with light-responsiveness, so that scanning a laser of an appropriate wavelength can displace the material, thus producing microtopographic features on the material surface (Bian et al. 1999). Photopatterning such materials with different microtopographies can be very effective in driving cellular morphology and migration (Rianna et al. 2015; Chiara Fedele et al. 2020). More interestingly, the microtopography can be reconfigured *in situ*, (Rianna et al. 2016), therefore paving the way for advanced studies of cellular response to ECM dynamics.

In addition, cellular responses occur in different time scales. For example, changes in cell shape, migration, and gene expression occur in a slow time scale of minutes to hours (Hoffman, Grashoff, and Schwartz 2011). However, many of these changes are triggered by initial, sub-second mechanotransduction processes, where physical cues are converted to biochemical activity. Here, mechanosensitive (MS) ion channels play a critical role as transmembrane proteins that are sensitive to mechanical forces and respond to mechanical stimulation in the millisecond-scale (Coste et al. 2010; Douguet and Honoré 2019). MS channels localize to the cell membrane and intracellular organelles, and they are gated so that they open due to e.g., increased membrane tension, allowing the transport of ions through them. Changes in local ion concentrations can then lead to subsequent changes in cellular signaling and functionality (Douguet and Honoré 2019).

For an array of physiological processes, Ca^2+^-mediating cation channels, including the mechanosensitive Piezo1 channel, are particularly important due to the dual function of Ca^2+^. In addition to affecting the cell membrane potential, Ca^2+^ acts as a universal second messenger with its immense concentration gradient over the cell membrane. Unlike any other ion, Ca^2+^ participates in numerous cellular signaling pathways, such as contraction, proliferation, secretion, vesicle trafficking, protein synthesis, and apoptosis (Clapham 2007; M. J. Berridge, Lipp, and Bootman 2000). Interestingly, increase in cytosolic Ca^2+^ concentration is often one of the first events in mechanotransduction (F. Li et al. 2018; Godin et al. 2007). Ca^2+^ signaling cascade is typically initiated by specific stimuli that result in the release of Ca^2+^ into the cytosol through ion channels at the cell membrane or endoplasmic/sarcoplasmic reticulum (ER/SR) membrane (M. J. Berridge, Lipp, and Bootman 2000; Clapham 2007). After the usually rapid rise in cytosolic Ca^2+^ concentration up to 1000 nM, the signal is turned off by various Ca^2+^ pumps and exchangers that restore the physiological cytosolic Ca^2+^ concentration down to 100 nM (M. J. Berridge, Lipp, and Bootman 2000; Clapham 2007). Ca^2+^ signals are spatiotemporally highly dynamic and can vary from local signals in proximity of the opening channel to spreading throughout the entire cell, and the durations of these signals can range from microseconds to hours (Boulware and Marchant 2008). Therefore, utilizing Ca^2+^ indicators to image Ca^2+^ dynamics offers a unique opportunity to observe the onset of mechanotransduction events. Furthermore, Ca^2+^ is an important intercellular messenger that allows tissue-wide communication (M. Berridge, Lipp, and Bootman 1999). Ca^2+^ signals can spread directly from cell to cell via gap junctions, transmembrane proteins that form pores interconnecting two adjacent cells (M. J. Berridge, Lipp, and Bootman 2000). This rapid gap junction mediated signaling can be amplified by Ca^2+^ induced Ca^2+^ release (CICR) (Roderick, Berridge, and Bootman 2003). Alternatively, Ca^2+^ signals can be distributed indirectly from cell to cell through slower pathways involving adenosine triphosphate (ATP) or inositol 1,4,5-trisphosphate (IP3) (M. J. Berridge, Lipp, and Bootman 2000). Therefore, Ca^2+^ also offers insight into the mechanically induced cell to cell communication.

Herein, we exploit the dynamic photopatterning of an azobenzene-based molecular glass (disperse red 1 molecular glass – DR1-glass (Kirby et al. 2014)) as a dynamic cell substrate for the light-induced mechanically activated observation of Ca^2+^ dynamics across an epithelial monolayer. We first optimize the light-induced formation of mechano-/topographical features by tuning irradiation parameters of a laser scanning confocal microscope (LSCM). We show that, in response to the shear deformation operated by local nanoscale modifications at the material surface, changes in cytosolic Ca^2+^ concentration are induced in the epithelium. Based on pharmacological channel modulation, we propose Piezo1 channels to be involved in sensing this deformation. Finally, this approach together with computational modeling reveals that cells respond by Ca^2+^ to direct mechanical stimuli and the subcellular mechanical stimulus is then spread to the neighbor cells. This enlightens the role of cell-to-cell communication in the tissues’ complex mechanoresponses.

## Results

### Photopatterning by scanning laser produces microtopographic features and material flow

The light-responsive thin films were prepared by spin coating DR1-glass on microscopy coverslips. The layer thickness (≈250 nm) was chosen so that it would not block the fluorescence from the used genetically encoded Ca^2+^ indicators expressed by the cells. The free-form microtopography was inscribed on the thin films by scanning a 488 nm laser by the galvanometric mirrors of a LSCM with a 63x/1.2 water-immersion objective. The laser was scanned only in user-defined regions of interest (ROIs) of the film, thus producing a microtopography on the film surface (Figure 1A, B), as reported previously by Rianna and co-workers on a different DR1-containing material (Rianna et al. 2016).

**Figure 1.**
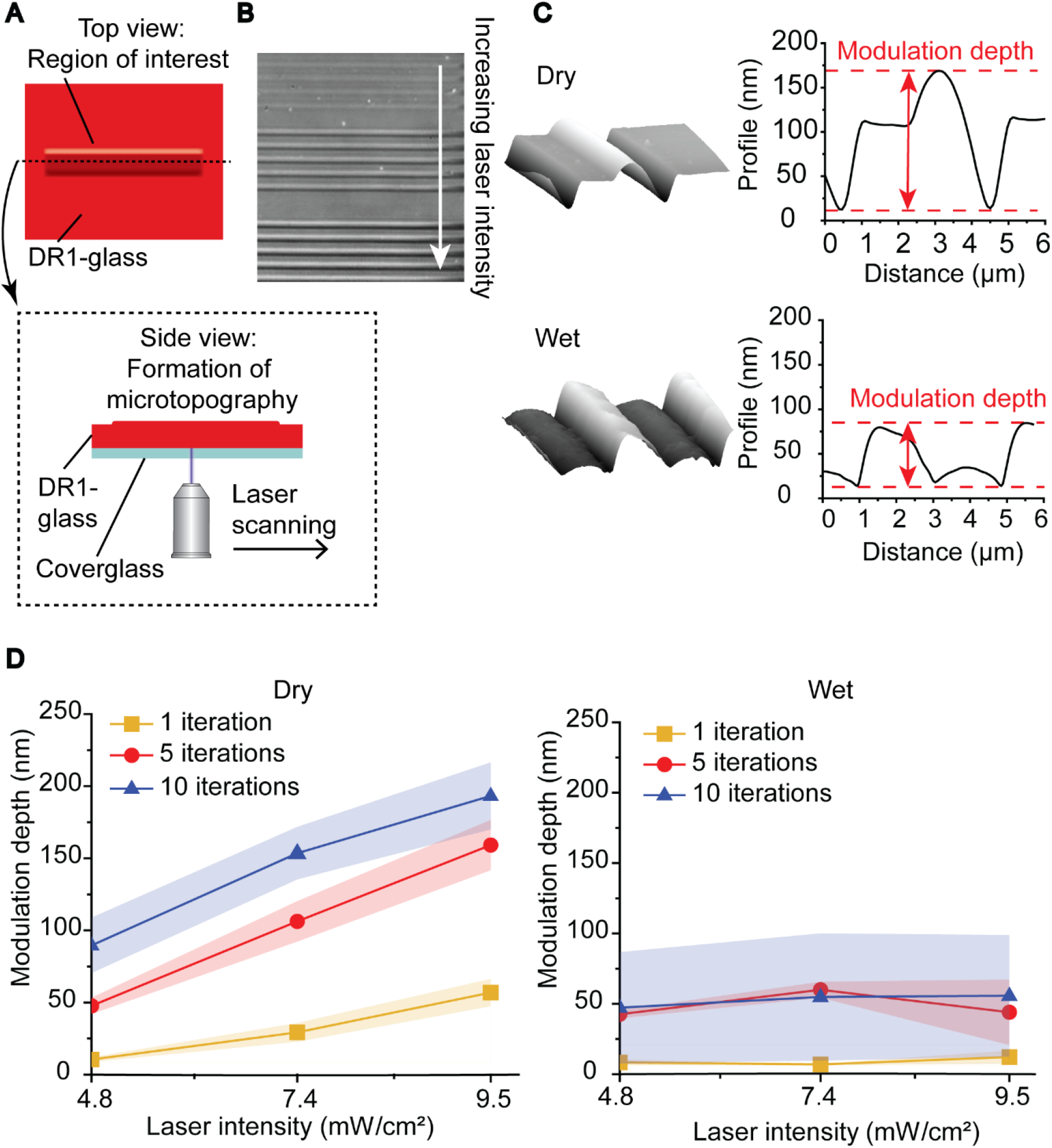
Characterization of light-induced microtopographies. A) Graphical representation of the laser scanning experiment. B) Bright field image of the effect of increasing intensity (from top to bottom) on the formed microtopography. C) AFM 3D projections of microtopography and cross-sectional profiles in dry and wet (7.4 mW/cm^2^, 10 iterations). D) Modulation depth of microtopographies as a function of laser intensity and number of iterations in dry and wet conditions.

We performed confocal patterning both in the absence of any medium (referred to as “dry”) and in the presence of a water-based medium (referred to as “wet”) to mimic the conditions necessary during cell culturing. The photoinscription parameters tested were the laser intensity (4.8 mW/cm^2^, 7.4 mW/cm^2^, and 9.5 mW/cm^2^), measured at the back focal plane of the objective (approximately corresponding to 3.7 J/cm^2^, 5.7 J/cm^2^, and 7.3 J/cm^2^ pixel radiant exposures at the objective focal plane; Figure S1A illustrates the measurements of the full laser intensity range) and the iteration cycles of the laser scanning process. Unless stated explicitly, the default setup for the laser scanning results in defined square pixel size (0.2 μm lateral size), pixel dwell time of 1 μs, and unidirectional laser scanning (i.e., laser scanning is performed lines by lines, from left to right). The two parameters that were varied during the experiments, the laser intensity and the number of iterations, determine the total energy received by each pixel of the material over the scanning:

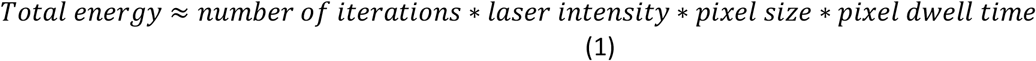

By increasing the laser intensity, the photon flux increases, whereas by increasing the number of iterations of the scanning process the exposure increases while keeping the photon flux constant. Following photoirradiation, the resulting microtopographic features were analyzed by atomic force microscopy (AFM) to evaluate the cross-sectional profiles (Figure 1C). The first evident difference between the inscribed features in dry and wet conditions is in the shape of the cross-sectional profiles. The dry environment favored the accumulation of the material in a well-defined dome-shaped line, whereas the presence of media on top of DR1-glass produced a flatter profile with smaller modulation depth, defined as the distance between the trough and the crest of the profiles. Furthermore, we noticed the emergence of a “spiky” texture at high radiant exposures (Figure S1B) that hampers the direct comparison of the topographical parameters identified. Such power settings were considered too high for the experiments due to photobleaching and possible phototoxicity to the cells and were eliminated from the parameters set.

Digital holographic microscopy (DHM) was used to measure the modulation depths over larger areas of the samples in all the replicates (Figure 1D). The average modulation depth was increasing with both laser intensity and number of iterations in dry conditions, whereas in wet environment it seemed generally independent of the laser intensity. The modulation depth can be discussed also in terms of the relative energy dose (Eq. 1 normalized to the first value and Figure S1C, D). The first observation was that for irradiation conditions that have the same relative energy dose, the irradiation in dry yields distinct modulation depths, with the deeper topography being obtained at higher laser intensity. On the contrary, in wet conditions photopatterning performed at the same relative energy yields topographies with the same modulation depth. Furthermore, in wet conditions we observed clear inhomogeneities in the shape of the features at higher relative energies, leading into higher deviation in the datapoints (Figure S1D).

We also studied the photo-induced modifications of the film in terms of their formation dynamics. The material displacement during laser scanning in dry conditions was characterized via Particle Image Velocimetry (PIV) (Thielicke and Sonntag 2021) through automatic tracing and cross-correlation of fluorescent particles deposited over the film surface (see Materials & Methods). Images were taken first in the transmission channel of the LSCM in between each iteration of the inscription process in dry conditions (Figure 2A). The PIV analysis confirmed that, upon laser scanning, DR1-glass moves in the direction opposite to the scanning direction, accumulating at the left side of the ROI (Supporting Video 1). Furthermore, the particles’ speed, calculated as the mean displacement of all particles inside the ROI after each frame, was constant during each iteration of the experiment (Figure 2B). The speed of this constant flow was dependent on the laser intensity as higher laser intensities led to faster material flow.

**Figure 2.**
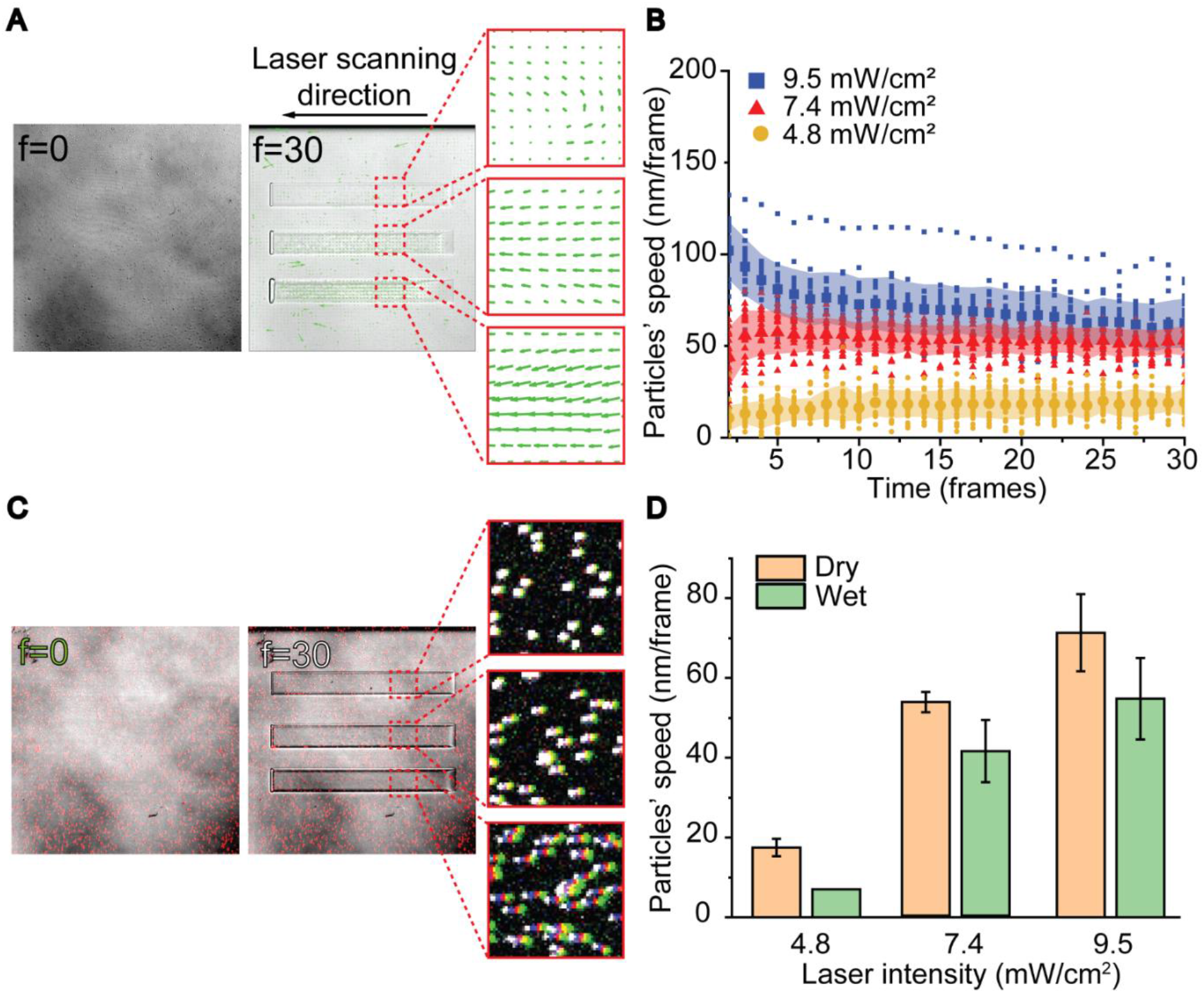
Characterization of material displacement. A) Overlay of bright field microscope images and PIV vector fields showing the material displacement at frame zero and frame 30. B) Particles’ speed as a function of time. C) Overlay of fluorescence images of particles at the initial (white) and final (green) frames and overlay of the entire time lapse to visualize the particles’ trajectories at different laser intensities (from top to bottom: 4.8, 7.4, and 9.5 mW/cm^2^. D) Mean particles’ speed during the photostimulation in dry (beige) and wet (green) conditions.

When performing the same experiment in wet conditions, we deposited a fibronectin coating onto the DR1-glass film and measured the particles’ displacement by comparing fluorescence images before and after the light stimulation (Figure 2C). The data indicated that also in wet conditions the speed of material displacement depended on the laser intensity, but the process was overall slower than in dry conditions (Figure 2D).

Thus far, we had used a unidirectional scanning mode, where the laser scans from left to right only. This results in piling of material in one end and in loss of material in the other end of the inscription (Figure 2A). Instead, with the bidirectional scanning mode, the laser scans each line of the ROI in zig-zag motion in both directions and therefore results in equal material accumulation on both sides of the stimulation region (Figure S2 and Supporting Videos 1 and 2). Despite the different macroscopic material movements, the achieved modulation depths are similar with both scanning modes in wet conditions (Figure S2).

In summary, we observed a clear difference in material behavior in wet and dry conditions, as in dry condition the topographies were higher (max ≈ 220 nm) and material flow faster (max ≈ 80 nm/iteration). Furthermore, in dry conditions, the modulation depth was affected by both the number of iterations and laser intensity. Interestingly, in the presence of water, which is fundamental for cell culturing experiments, we noticed that the modulation height of the topography (max ≈ 100 nm) was dependent on the number of iterations and independent of laser intensity, whereas the speed of lateral displacement (max ≈ 65 nm/iteration) was determined by laser intensity. Fluorescent particle tracking showed that these lateral displacements are reproducible for every laser scanning iteration (up to 30) of DR1-glass photostimulation (Figure 2B). This means that with this technique we can apply shear forces to cells by simply modulating the laser intensity. This approach is appealing because cells can be restimulated mechanically in arbitrary intervals after the initial round of photostimulation. Added to this, these rounds of stimulation can also be applied by using any pattern size or shape, thus allowing for localized as well as tissue-level mechanical stimulation. This degree of freedom in spatiotemporal control of the mechanical perturbations revealed here will be instrumental in studying several processes of different time scales in cellular responses and how cells in tissues collectively respond to the ECM dynamics.

### Lateral displacement of the DR1-glass layer induces intracellular Ca^2+^ responses in MDCK II cells

Previous experiments indicated that the magnitude and dynamics of the photoinduced topographical changes in the DR1-glass can be precisely controlled by tuning the laser scanning speed and iterations. Since the substrate is biocompatible, cells can be cultured on the material, which allows the studies of cellular responses to nanoscale topographical movements in the cell-ECM interface. The focused laser photoinscription occurring in microsecond timescale can act as programmable mechanical stimulator of the cells. To investigate this, cells constitutively expressing red-emitting (emission maxima approx. 605 nm) genetically encoded Ca^2+^ indicator (GECI) jRCaMP1b (Dana et al. 2016) were cultured on fibronectin-coated DR1-glass. Due to the minimal overlap in the excitation-emission spectra of the Ca^2+^ indicator (excitation maxima approx. 560 nm) and DR1-glass (absorption maxima approx. 480 nm) (Figure S3), the imaging of the Ca^2+^ indicator and photostimulation of the 250 nm - thick DR1-glass could be conducted sequentially. Via time lapse imaging using LSCM (Figure 3A), we monitored the dynamics of cytosolic Ca^2+^ concentration in single cells via following the jRCaMP1b fluorescence. Prior to photostimulation, we could observe a stable baseline level of cytosolic Ca^2+^, with few cells exhibiting low-amplitude spontaneous fluctuations in Ca^2+^ concentration characteristic to MDCK cells (Udagawa et al. 2012).

**Figure 3.**
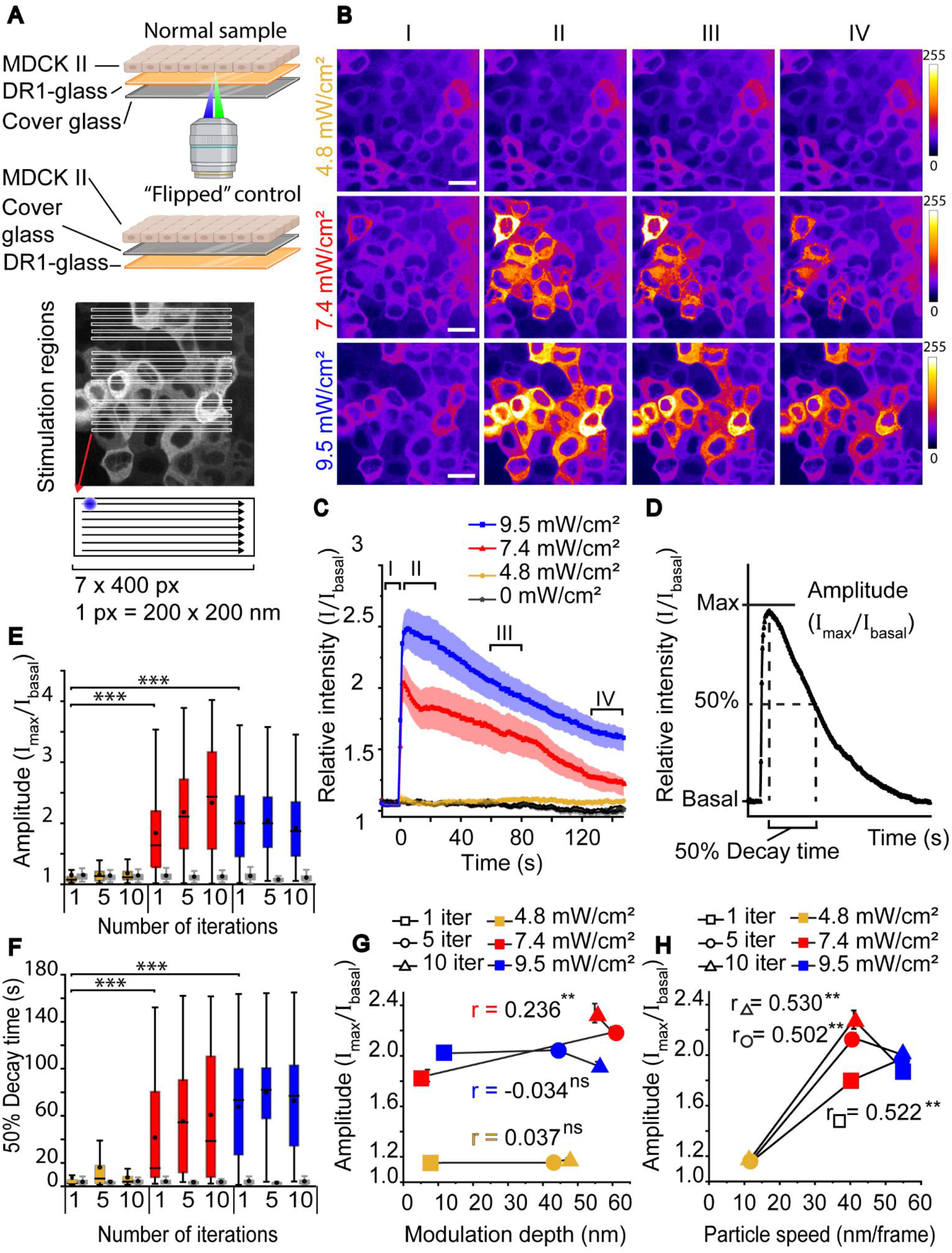
Characterization of cell responses. **A)** Top: A schematic representation of a sample with cells cultured on top of a fibronectin-coated DR1-glass layer, and a “flipped” control where the DR1-glass layer is positioned beneath the cover glass and therefore there is no interface between cells and the DR1-glass. Imaging (green laser beam) and DR1-glass stimulation (blue laser beam) is performed from below the sample. Below: The stimulation regions consisted of 3 x 5 rectangles á 7 x 400 pixels. The blue spot marks the size of the laser beam. Only cells in direct contact with the regions were analyzed. **B)** Average intensity projections of cells stimulated with 4.8 mW/cm^2^ (yellow), 7.4 mW/cm^2^ (red), and 9.5 mW/cm^2^ (blue) laser intensity before stimulation (I), during first 25 s after stimulation (II) and during the decay period (III 60-85 s after stimulation and IV 120-145 s after stimulation). Scale bar 20 μm. Color bar represents calcium signal intensity. **C)** Mean ± SE intensity plots of cells stimulated as in (B) and without stimulation (grey). The time intervals I-IV are marked in the plot. **D)** The maximum relative signal intensity is referred to as amplitude, and the time from max intensity to 50% max intensity is referred to as the 50% decay time. **E)** Amplitudes and **F)** 50% decay times of cells stimulated with 1, 5 or 10 iterations with 4.8 mW/cm^2^, 7.4 mW/cm^2^, and 9.5 mW/cm^2^ laser intensity. Plots marked in grey are for the corresponding flipped control samples. In box plots the boxes mark the 25% - 75% interquartile ranges (IQR), whiskers mark the range within 1.5 x IQR, horizontal lines mark the median and circles mark the mean. Statistical significances were calculated with Kruskal-Wallis test. **G)** Amplitudes as a function of modulation depth with Pearson’s correlations (r). Weak or no correlation is detected. **H)** Amplitudes as a function of particle speed with Pearson’s correlations (r). A strong correlation is detected between the speed of flow and the resulting cell response. **E-H)** For normal samples the sample size varies from n = 138 to n = 639 and for flipped control samples from n = 77 to n = 278. Statistical significances are shown as ns = p-value > 0.05, * = p-value ≤ 0.05, ** = p-value ≤ 0.01 and *** = p-value ≤ 0.001.

We photopatterned the DR1-glass with unidirectional laser scanning in a ROI containing 15 stripe-like regions of 7 x 400 pixels (1.4 μm x 80 μm) (Figure 3A). This setting is referred to as the multi-region setting, in contrast to single-region setting used in the next sections, where ROI contains only 1 stripe-like region. We stimulated the DR1-glass with different irradiation parameters (4.8 mW/cm^2^, 7.4 mW/cm^2^, and 9.5 mW/cm^2^ laser intensity with 1, 5 and 10 iterations) and recorded the cellular Ca^2+^ responses via the jRCamP1b signals (Figure 3B). The signal in each cell was normalized to the initial sensor intensity in the corresponding cell to account for variability in the constitutively expressed jRCamP1b protein and basal cytosolic Ca^2+^ level. By plotting normalized Ca^2+^ responses against time (Figure 3C), we observed the typical response to exhibit a Ca^2+^ “surge” that begins immediately after stimulation and lasts for 2-5 seconds, followed by a slow decay to the baseline level. Here, we only analyzed cells that were directly perturbed by mechanical stimulations, i.e., were in contact with the stimulation region, although Ca^2+^ responses were detectable also in some cells further away.

We extracted two key features from the normalized Ca^2+^ trace in each cell: the amplitude, which is the maximum intensity relative to the basal intensity, and the 50% decay time, which we determined as the time it takes for the signal to decrease from the maximum down to 50 % of the maximum (Figure 3D). Figures 3F-G show that the cell responses to 4.8 mW/cm^2^ laser intensity stimulation have negligible amplitudes and short decay times, with few cells showing intensive responses either to 1, 5 and 10 iterations. With 7.4 mW/cm^2^ and 9.5 mW/cm^2^ laser intensities, the amplitudes show similar, substantial increase from the baseline intensity (Figure 3E). Despite having similar amplitudes, their 50% decay times, while both being longer than that with 4.8 mW/cm^2^ laser intensity, differ: the decay time with 9.5 mW/cm^2^ laser intensity are ~30 s longer than that with 7.4 mW/cm^2^ (Figure 3F). Notably, increasing the number of photoirradiation iterations slightly increases the amplitudes and decay time with 7.4 mW/cm^2^ but with 9.5 mW/cm^2^ laser intensity, no such increase is seen.

In addition to the apparent differences in population-wide Ca^2+^ responses with different laser intensities (Figure 3E-F), we also detected high level of variability in Ca^2+^ dynamics among individual cells. Even with 7.4 mW/cm^2^ and 9.5 mW/cm^2^ laser intensities, which showed clear responses in population level, only part of the stimulated cells (ROI shown in Figure 3A) show a response while others remain quiet (Figure 3B). This may be due to the cell-to-cell heterogeneity in the molecular components involved in the Ca^2+^ responses. Such heterogeneity has been reported for example in cell responses to stretch (Latorre et al. 2018) and compression (Peussa et al. 2022). Also, MDCK II cells cultured as monolayers, when matured, form fluid-filled dome structures into the monolayer (Choudhury et al. 2022), which may render some cells unreachable by photostimulation.

To verify that the observed Ca^2+^ responses were induced by topographical changes in the DR1-glass and not due to phototoxic effects of the stimulating laser, we performed identical stimulation experiments with “flipped” control samples (Figure 3A), where the DR1-glass coating and cell culture were placed on different sides of the fibronectin-coated cover glass. In flipped samples, cells experienced no mechanical stimulation, but the presence of DR1-glass guaranteed similar absorption conditions, thus allowing us to detect the effects of phototoxicity. Indeed, no Ca^2+^ responses were detected in the flipped samples (Figure 3E-F, grey boxes) with any stimulation parameters. We stimulated flipped samples also with intensities beyond the presented data (up to 14.3 mW/cm^2^) but saw no cell responses.

We also investigated whether the possible generation of heat from DR1-glass photostimulation, rather than material displacements, can trigger cell responses. Transient local heat may be generated by increasing intensity of the photostimulation and it may accumulate or be efficiently dissipated, depending on the heat capacity of the surrounding medium (Yager and Barrett 2003). We hypothesized that, by increasing the exposure, heat accumulation would occur in the irradiated spots. Thus, as opposed to directly increasing the laser intensity, we would be able to generate a cell response simply by increasing either the number of iterations or the dwell time of the laser scanning. To test this, we used the lowest laser intensity (4.8 mW/cm^2^), which did not produce Ca^2+^ responses with 1 μs pixel dwell time, and increased the dwell time to 2.55 μs, 12.6 μs and finally 177 μs (Figure S3). With 2.55 μs dwell time the 4.8 mW/cm^2^ laser intensity leads to similar relative energy as achieved with 1 μs dwell time using 7.4 mW/cm^2^ and 9.5 mW/cm^2^ laser intensities, and with 12.6 μs the exposure is already ten-fold larger. Yet, no cell responses were detected. Only with 177 μs dwell time, which leads to a substantially higher exposure than what we used in our other experiments, we saw a rise in Ca^2+^ signals. This suggests that within the range of exposure that we applied, heat accumulation was insignificant. Furthermore, the presence of water may efficiently dissipate some of the possibly generated heat, thus decreasing the thermal effects on cells. In addition, at high laser intensities, photothermal effects in DR1-containing materials may negatively affect the photopatterning efficiency when the material locally reaches its glass transition temperature (Tofini et al. 2018). We did not observe any decrease in modulation depth with increasing laser intensity (Figure 1D and Figure S1), which suggests that such temperatures were not reached, not even in dry conditions. Thus, even if we cannot completely rule out thermal effects, our data suggests that the dominating cellular stimulus is mechanical.

Since photostimulation modulates the DR1-glass in two different ways, i.e., the formation of vertical topography (Figure 1) and lateral displacement (Figure 2), we sought to discover which of these movements trigger the Ca^2+^ responses in cells. We plotted amplitudes of cell responses against resulting modulation depth (Figure 3G) and lateral particle speed (Figure 3H). We found Spearman correlation between amplitude and modulation depth to be weak at best (r<0.24**) (Figure 3G). Thus, the cellular responses did not strongly depend on the height of the resulting topographical features. On the other hand, we found a strong positive correlation between the lateral particle speed and the amplitude (r>0.5** for each number of iterations, Figure 3H). This indicated that the cells responded mainly to shear deformation resulting from the lateral flow of the DR1-glass. Additionally, 10 iterations with 4.8 mW/cm^2^ results in similar total deformation (Figure 2D) as 1 iteration with 7.4 mW/cm^2^, but cell responses were only detected with the higher laser intensity. This indicates that the speed of the deformation, not only magnitude of the total deformation, was crucial.

To determine the scope of mechanical stimulation that cells respond to, we repeated the experiments with the bidirectional laser scanning mode. With unidirectional laser scanning, the material in the ROI flows laterally in a single direction (Figure S2), which should exert macroscopic basal shears to cells and potentially deform them as cells in ROI are displaced. With bidirectional laser scanning, no apparent lateral flow was observed but anisotropic displacements of the material are visible with LSCM (Figure S2). Thus, shear, if any, should only occur at the sub-micron levels. Interestingly, changing the scanning mode did not affect the Ca^2+^ signal amplitude, and only caused a small increase in the decay time (Figure S3). This suggests that very local, sub-micron movements in the cell-material interface were leading to the cellular response.

Altogether, we found that the photoinscribed dynamic topographical changes in the DR1-glass layer can trigger mechanically induced Ca^2+^ signals in epithelial cells. We determined that the cell responses detected here are triggered by shear deformation, caused by the material lateral flow, instead of the modulation depth of the microtopography. Cell responses were triggered by as slow as 1.48 nm/ms shear displacements (41 nm/frame particle speed achieved with 7.4 mW/cm^2^ laser intensity, and the inscription time per one stripe is 27.7 ms). Furthermore, we showed that the cell responses were triggered by very local, sub-micron material displacements, instead of macroscopic net flow of material.

### Ca^2+^ signals are triggered by mechanosensing Piezo1 channels and amplified in the ER

Next, we investigated the mechanism responsible for the detected Ca^2+^ responses. The Ca^2+^ surges in response to shear deformation were very fast, occurring immediately after the mechanical stimulation (Figure 3B-C). The speed of the responses pointed toward MS ion channels, the fastest responders to mechanical stimulations (Coste et al. 2010; Douguet and Honoré 2019). Since the cells are adhered to the surface via integrin-rich adhesions (Martino et al. 2018; Iskratsch, Wolfenson, and Sheetz 2014), nanoscale material movements can lead to increased local tensions in the cell membranes. MDCK cells express mechanosensitive Piezo1 channels (Gudipaty et al. 2017), which have been found to localize also at the basal cell membrane (Caceres et al. 2019). Moreover, Piezo1 channels are sensitive to changes in the membrane tension, and can induce Ca^2+^ signals in epithelial cells (Gudipaty et al. 2017; Jetta et al. 2019). Therefore, we hypothesized that the material deformation could generate tension in the cell membrane, leading to the opening of Piezo1 channels and subsequent Ca^2+^ influx into the cell. Thus, we immunostained Piezo1 channels and used confocal microscopy to determine its intracellular localization. The imaging indicated that the Piezo1 channels are abundantly expressed in MDCK II cells (Figure 4A). Furthermore, they localized to the basal side of the cells, similarly to the actin stress fibers (Figure 4B), suggesting that the channels are indeed located close to the cell-DR1-glass interface. This supports the hypothesis that Piezo1 channels could be responsible for the detected cell responses.

**Figure 4.**
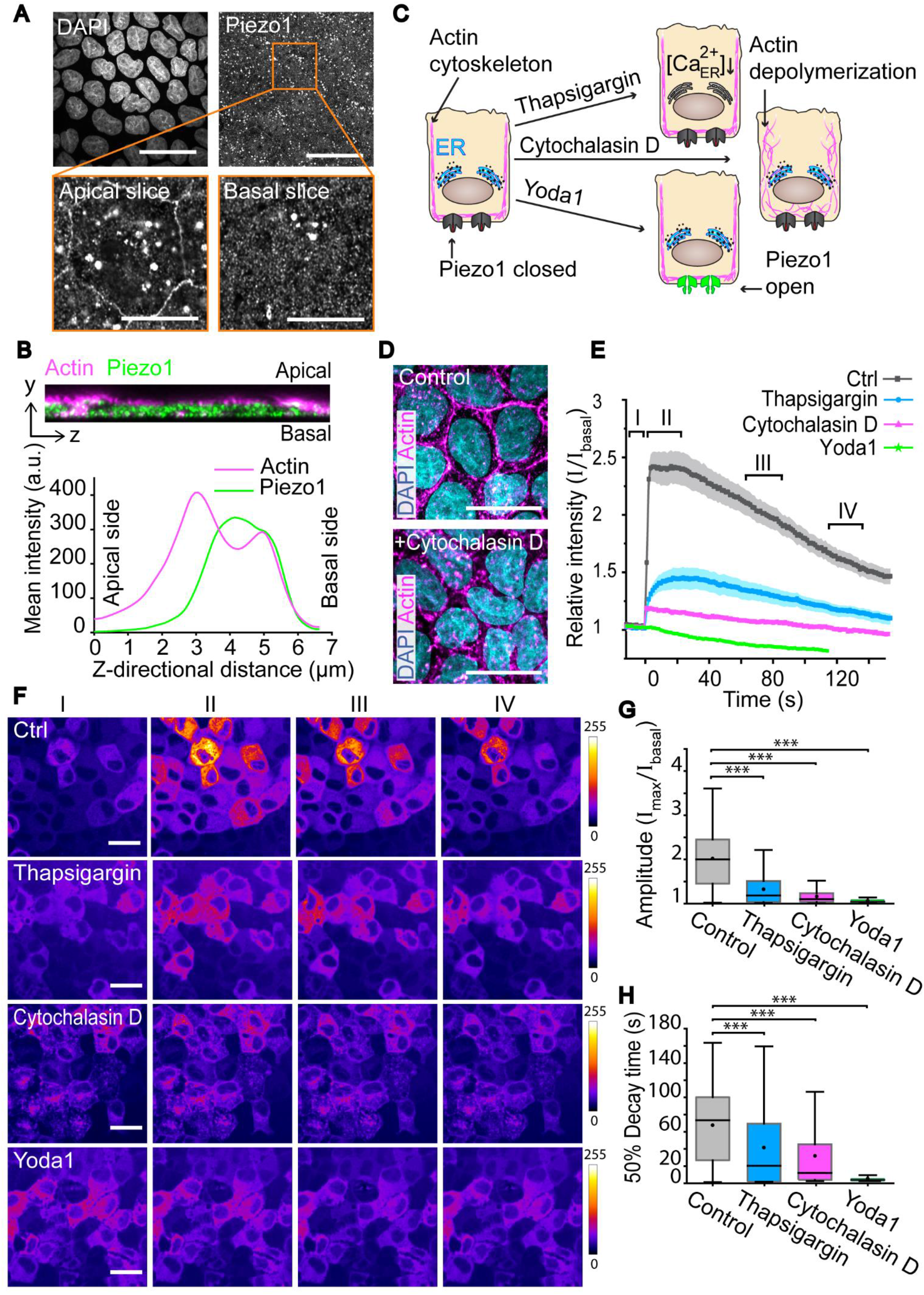
Effects of Cytochalasin D, thapsigargin and Yoda1 on cell responses. **A)** Maximum intensity projections showing the nuclei (DAPI), and Piezo1. Scale bars 30 μm. Below, magnification (orange box) displaying single slices of the Z-stack from the apical and basal membranes showing the Piezo1 channels. Scale bars 10 μm. **B)** A 20-slice orthogonal Z-projection and the mean distribution of actin (magenta) and Piezo1 (green) signal intensity measured from the magnified cell in (A)). **C)** Schematic representation of the effects of the drugs. Thapsigargin depletes intracellular Ca^2+^ stores, cytochalasin D inhibits the polymerization the actin cytoskeleton, and Yoda1 facilitates opening of Piezo1 channels. **D)** Cells without and after 1 h treatment with cytochalasin D showing nuclei (DAPI, cyan) and actins (phalloidin, magenta). Scale bars 20 μm. **E)** Mean ± SE intensity plots of cell responses in control conditions and after treatment with cytochalasin D, thapsigargin, or Yoda1. **F)** Average intensity Z-projections of cells before stimulation (I), during first 25 s after stimulation (II) and during the decay period (III 60-85 s after stimulation and IV 120-145 s after stimulation). Scale bars 20 μm. Color bar represents calcium signal intensity. **G)** Amplitudes and **H)** decay times of cell responses without drug, or after treatment with thapsigargin (n= 130), Cytochalasin D (n = 130) or Yoda1 (n = 99). In box plots the boxes mark the 25% - 75% interquartile ranges (IQR), whiskers mark the range within 1.5 x IQR, horizontal lines mark the median and circles mark the mean. Statistical significances (Kruskal-Wallis test) *** = p-value ≤ 0.001.

To test the hypothesis, we used two pharmaceutical drugs, Yoda1 and Cytochalasin D, which are known to affect Piezo1 activation either directly or indirectly (Figure 4C) and thus should affect the Ca^2+^ signals if influx occurs via Piezo1 channels. Cytochalasin D inhibits the polymerization of the actin cytoskeleton and therefore reduces the tension in cortical actin network and cell membrane. This indirectly affects Piezo1 activation (Casella, Flanagan, and Lin 1981) (Figure 4C). As Piezo1 channels open due to increased membrane tension, loosening the membrane disables the channels’ mechanosensing ability. To verify the effects of Cytochalasin D, we immunostained samples with phalloidin before and after Cytochalasin D treatment. One-hour treatment with 10 μg/ml Cytochalasin D strongly disrupted the actin cytoskeleton (Figure 4D). This treatment also significantly decreased the amplitude of Ca^2+^ signals (Figure 4E-G and Supporting Video 5), suggesting that the mechanical stimulation was less effectively transduced. Consequently, also the decay time decreased significantly (Figure 4H). We also used GsMTx-4, spider venom that locally relaxes the cell membrane around Piezo1 channels and therefore renders the channels less sensitive to forces (Gnanasambandam et al. 2017), but we saw no change in cell responses. We believe that the forces generated by our system may simply be so large that the effect of GsMTx-4 remained insignificant. However, the combined results from Yoda1 and Cytochalasin D experiments strongly suggest that Piezo1 channels play a key role in epithelial mechanosensation stimulated through basal shear. A possibility remains that other MS channels besides Piezo1s are also involved in the generation of the detected Ca^2+^ signals, but their involvement was not further studied here.

Next, we performed experiments with Yoda1, which decreases the activation threshold for Piezo1, making channels more sensitive to membrane tension, (Botello-Smith et al. 2019). After 10 μM addition of Yoda1 we saw a high and sustained increase in intracellular Ca^2+^ concentration. This verifies the presence and functioning of Piezo1 channels in MDCK II cells observed with immunostainings. Cells were allowed to relax to a new equilibrium before performing photostimulation to the DR1-glass layer. Interestingly, one iteration of photostimulation of DR1-glass with 9.5 mW/cm^2^ laser intensity performed after the adjustment period failed to produce any Ca^2+^ response in the cells (Figure 4E-G and Supporting Video 4). Although the baseline Ca^2+^ level of cells was elevated after Yoda1 treatment and demonstrated functional Piezo1 channels, the lack of any Ca^2+^ response further corroborated the involvement of Piezo1 channels: once all Piezo1 channels had been forced open with Yoda1, the mechanical stimulation could no longer produce an additional Ca^2+^ influx via them.

In addition to ion channels at the cell membrane, Ca^2+^ signaling involves the participation of other Ca^2+^ transport proteins, especially at the sarco-endoplasmic reticulum (SR/ER) (M. J. Berridge, Lipp, and Bootman 2000; Clapham 2007). To probe the role of intracellular Ca^2+^ stores in the recorded Ca^2+^ responses, we performed experiments with thapsigargin, a potent SR/ER calcium-ATPase (SERCA) inhibitor, which causes depletion of the intracellular Ca^2+^ stores (Thastrup et al. 1990) (Figure 4C). Thapsigargin (1 μM) was added to the cells and after 15 min relaxation period the cells were mechanically stimulated via DR1-glass photoinscription. We saw a significant decrease, but not complete elimination, of Ca^2+^ response amplitudes once the intracellular Ca^2+^ stores had been depleted (Figure 4G and Supporting Video 6). The 50% decay time was likewise shortened (Figure 4H). This suggests that, in normal conditions, additional Ca^2+^ is released from the ER following the initial Ca^2+^ influx via Piezo1 channels, typical to Ca^2+^-induced Ca^2+^ release. Therefore, the cytosolic Ca^2+^ surge observed is a combination of influx from extracellular and intracellular sources.

Our data suggests that Piezo1 channels on the basal membrane are involved in sensing the DR1-glass generated material lateral flow. Opening of the Piezo1 channels permits an influx of extracellular Ca^2+^ and possibly other cations, which lead to rise in intracellular Ca^2+^ concentration. In addition, the signal is further amplified from internal Ca^2+^ stores most likely via CICR. However, the used imaging speed (1 frame per second) did not allow us to distinguish between the Ca^2+^ influx through Piezo1 channels and the positive feedback from CICR. Combined with data in the previous sections, our findings shed light on the mechanical cell-ECM interactions and on Piezo1 channels. We found that the Piezo1-mediated Ca^2+^ response can be achieved with a material displacement as small as 40 nm, which translates to 1.48 nm/ms (40 nm/frame particle speed achieved with 7.8 mW/cm^2^ laser intensity, and the inscription time per one rectangle is 27.7 ms). This is in the same range as the 10 nm displacement reported by Poole et al (Poole et al. 2014) to trigger Piezo1 channels’ opening, thus giving more evidence on the sensitivity range of Piezo1 channels. Our data also concurs with previous findings showing that Piezo1 channels are located on the basal membrane (Caceres et al. 2019; Ellefsen et al. 2019). Furthermore, we were able to pinpoint that in our system Piezo1 channels specifically sensed forces parallel to the cell membrane i.e., basal shear forces rising from the material movement. Apical shear forces are known to be present for example in endothelium from blood and lymph flow, and Piezo1 channels have been shown to sense apical shear (Jetta et al. 2019), but there has not been much focus on shear forces on the basal side.

Basal mechanical cell-ECM interactions are known to play a role in, for example, cancer progression and in development, but the details of this mechanical coupling are mostly unknown. It has been shown that the stiffness of the basement membrane, a specialized type of ECM that forms a boundary between different tissue types, affects cancer metastasis (Reuten et al. 2021), and that cancer cells can cleave the ECM by secreting metalloproteases (Niland, Riscanevo, and Eble 2021) thus changing the force field in the ECM. Dynamic changes in ECM density are also related to epithelial fold formation during development (Sui et al. 2018). In recent work, Ellefsen et al. (Ellefsen et al. 2019) showed that Piezo1 channels are localized evenly throughout the basal membrane but show most activity near focal adhesions. With this, we hypothesize that focal adhesions may generate also shear forces to their proximity, thus activating Piezo1 channels. Based on our findings, we propose that shear forces may play an unexpected part in basal cell-ECM interactions, for example, at focal adhesions. The proposed materials concept is promising for studying such effects in more detail.

### Ca^2+^ response kinetics differ in stimulated and in neighbor cells

In the previous experiments, we noticed that Ca^2+^ signals were not limited only to these cells in the photostimulated DR1 region. Instead, signals were also detected outside the stimulated area, signifying the involvement of intercellular communication (Figure 3B). To further explore this communication, we restricted DR1-glass stimulation to a single region of illumination (7×400 pixels, 1.4 μm x 80 μm), with 1 iteration of 9.5 mW/cm^2^ laser intensity, pixel dwell time of 1 μs (Figure 5A). In these single-region stimulation experiments (in contrast to the multi-region stimulation experiments above), we followed: i) *target cells* (purple) with overlapping surface contact area with the stimulation region; ii) *neighbor cells* (yellow) as adjacent cells of the target cells but not overlapping with the stimulation region; and iii) *others* (green) (Figure 5A). Here, with the imaging frame rate of ~1 frame/s, the Ca^2+^ signals in target cells and in neighbor cells occurred almost simultaneously (Figure 5A). We believe that a higher temporal resolution imaging may help distinguish the signal onset in target and neighbor cells but at the expense of imaging area. Nevertheless, this speed at which the responses spread pointed towards gap junction-mediated signaling (Frame and de Feijter 1997). Gap junctions are transmembrane proteins that form pores interconnecting two adjacent cells and thus enable Ca^2+^ signals to rapidly spread directly from cell to cell (M. J. Berridge, Lipp, and Bootman 2000), and gap junction based connectivity is known to exist in epithelial cells (Ponce et al. 2014; Mally et al. 2006).

**Figure 5.**
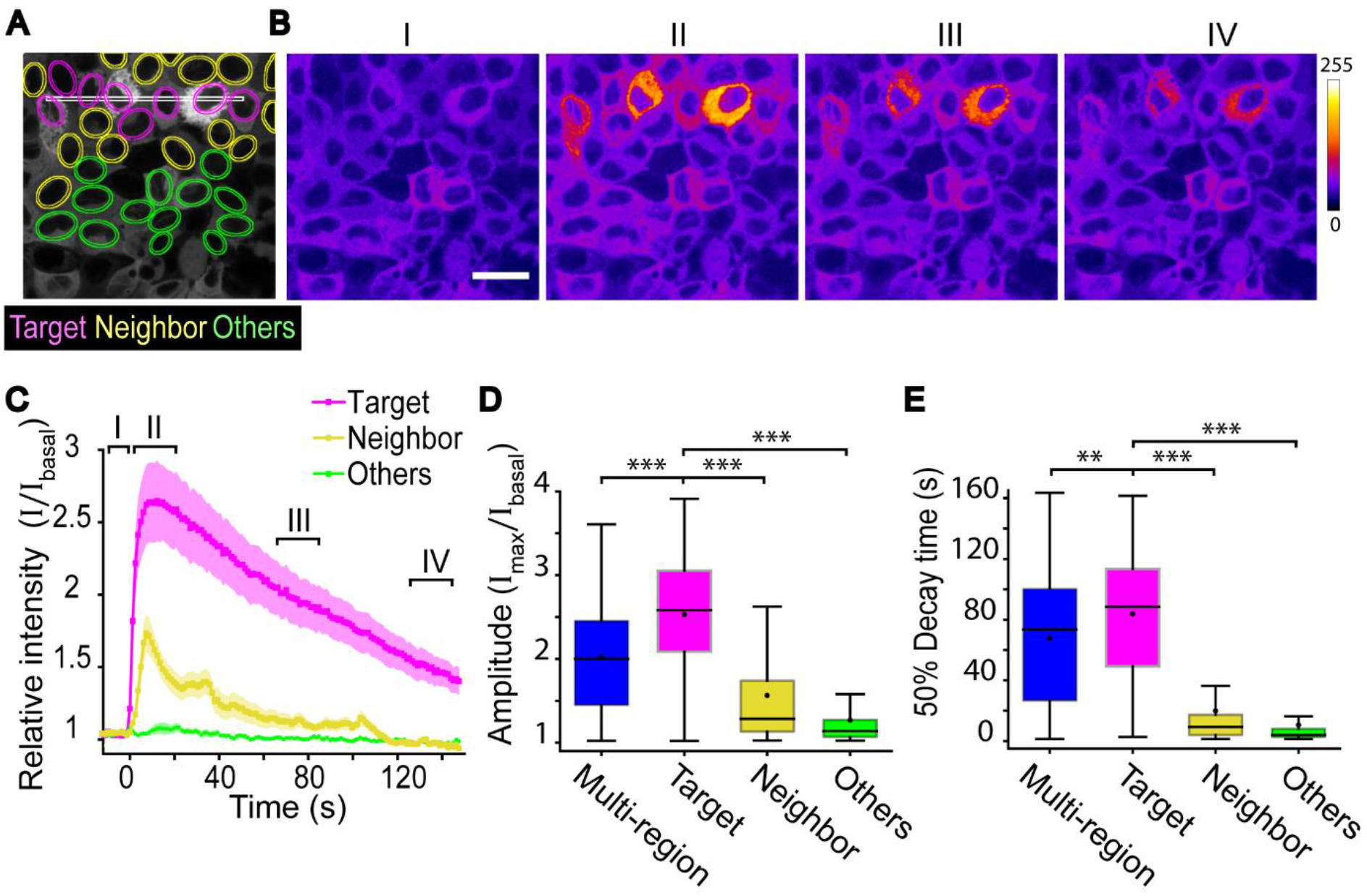
Propagation of Ca^2+^ signal within the epithelial monolayer. **A)** Representative stimulation experiment indicating Stimulation region (white), target cells (magenta) neighbors (yellow), and others (green). **B)** Average intensity Z-projections before stimulation (I), during first 25 s after stimulation (II) and during the decay period (III 60-85 s and IV 120-145 s). Scale bar 20 μm. Color bar represents calcium signal intensity. **C)** Mean ±SE of Ca^2+^ signal intensity over time in different cell groups presented in A with single-region photostimulation. **D)** Amplitudes and **E)** decay times of cells’ Ca^2+^ responses with either multi-region stimulation (blue, n = 639), or with single-region stimulation in target cells (purple, n = 183) in neighbor cells (yellow, n = 275) and in other cells (green, n = 472). Kruskal-Wallis test shows the statistical significance.^+^. **D-E)** Boxes mark the 25% - 75% interquartile ranges (IQR), whiskers mark the range within 1.5 x IQR, horizontal lines mark the median and circles mark the mean. Statistical significances are shown as ** = p-value ≤ 0.01 and *** = p-value ≤ 0.001.

Mechanically induced Ca^2+^ transients have also been observed in other epithelial cells, e.g., in retinal pigment epithelium (Abu Khamidakh et al. 2013). However, in these studies the stimulation was typically performed apically. Therefore, as a positive control experiment, we used micromanipulation to apically stimulate a single cell in the MDCK II epithelium (see Materials & Methods). Here, we also detected Ca^2+^ response in the targeted cell, in its neighbor cells and in further cells (Figure S4 and Supporting Video 7). This supports that Ca^2+^ signals detected outside the stimulated area are indeed triggered by intracellular communication and not by unintentional DR1-glass-stimulation. Notably, with apical stimulation the Ca^2+^ signal seemed to spread further than with DR1-glass stimulation. As the apical stimulation could not be as sophisticatedly controlled as DR1-glass stimulation, the triggering mechanical force may have been larger than in DR1-glass mediated stimulation, and thus could explain the different kinetics. Alternatively, this may suggest that apical and basal stimulation could activate different intercellular Ca^2+^ signaling processes. This would indicate that cells are capable of also spreading information about the direction of the mechanical cue. As the nature of stimulations (apical poking versus basal shear) were different, the cell responses were not further compared.

Next, we compared the DR1-glass evoked Ca^2+^ responses across the different cell groups. With single-region stimulation, we observed that Ca^2+^ responses in directly stimulated target cells had significantly higher amplitude and slower decay time than in the neighbor cells. In other more-distant cells, the signals were even smaller and less common (Figure 5D-E). When comparing two sub-populations of target and neighbor cells with similar amplitude distribution (see Figure S5A, with an example in Figure 5D), we verified that the difference in the decay time in these two populations was not simply due to the difference in amplitude, but that the cell groups exhibited different signaling kinetics.

Interestingly, we noticed that in single-region experiments the amplitudes of Ca^2+^ responses in targeted cells were, on average, larger than in multi-region experiments (Figure 5D) despite having same stimulation parameters (9.5 mW/cm^2^, 1 iteration). This is peculiar given that, in multi-region experiments, cells receive stimulation from a larger surface area and thus are expected to exhibit greater responses. The mean 50% decay time in multi-region experiments was also slightly shorter than that in target cells but longer than that in neighbor cells (Figure 5E). Thus, we hypothesize that the responses in multi-region experiments are a mixture of responses from direct mechanical stimulations (as in target cells) and from intercellular communication (as in neighbor cells). This mixture of responses may explain the higher variation detected in cell responses in multi-region experiment in comparison (Figure 5D-E). The finding further supports the theory that cell populations are not homogenous, but instead individual cells have differing tasks (Latorre et al. 2018; Peussa et al. 2022).

Next, we investigated the difference in the 50% decay time in target and neighbor cells. We verified that this difference is not simply caused by the difference in the response amplitude (Supporting Information Section A and Figure S5A), i.e., the initial influx of Ca^2+^ through the mechanically opened Piezo1-channels in target cells or through gap junctions in neighbor cells. Instead, the data suggested that there was a difference in the machinery responsible for restoring the baseline Ca^2+^ concentration. Although Piezo1-channels favor Ca^2+^, they are non-specific cation channels and thus also permit the flow of other cations, including Na^+^ (Romac et al. 2018). We therefore propose that in target cells the detected influx of Ca^2+^ is accompanied by Na^+^. This increase in cytosolic Na^+^ concentration could affect the efficiency of Na^+^/Ca^2+^ exchangers (NCX) (Dipolo and Beaugé 2006), one of the components responsible for removing cytosolic Ca^2+^. As NCXs use the Na^+^ gradient across cell membrane as an energy source, increasing Na^+^ concentration would decrease the pumping rate of Ca^2+^ out of the cells. Ultimately, this would result in a longer decay time in target cells. Meanwhile, in neighbor cells no Piezo1-mediated Na^+^ influx would occur. Unlike Ca^2+^ signals, Na^+^ signals is not amplified from the ER and thus would not suffice in generating significant Na^+^ influx to neighbor cells. Thus, we assumed that in neighbor cells the Na^+^ concentration remained unperturbed, and hence Ca^2+^ pumping rate out of cytosol remains high, leading to a shorter decay time as observed from the experiments (Figure 5C).

To test this hypothesis, we extended the simplified model of Ca^2+^ response in Kaouri et al. (Kaouri et al. 2019) to account for Na^+^ cytosolic concentration (Figure 6A). In this model, stimulations either from DR1-glass displacements or from adjacent cells trigger an indistinguishable Ca^2+^ influx *J_θ_* in target cells and neighbor cells. However, the influx of Na^+^ 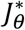 is set:

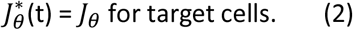

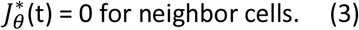

**Figure 6.**
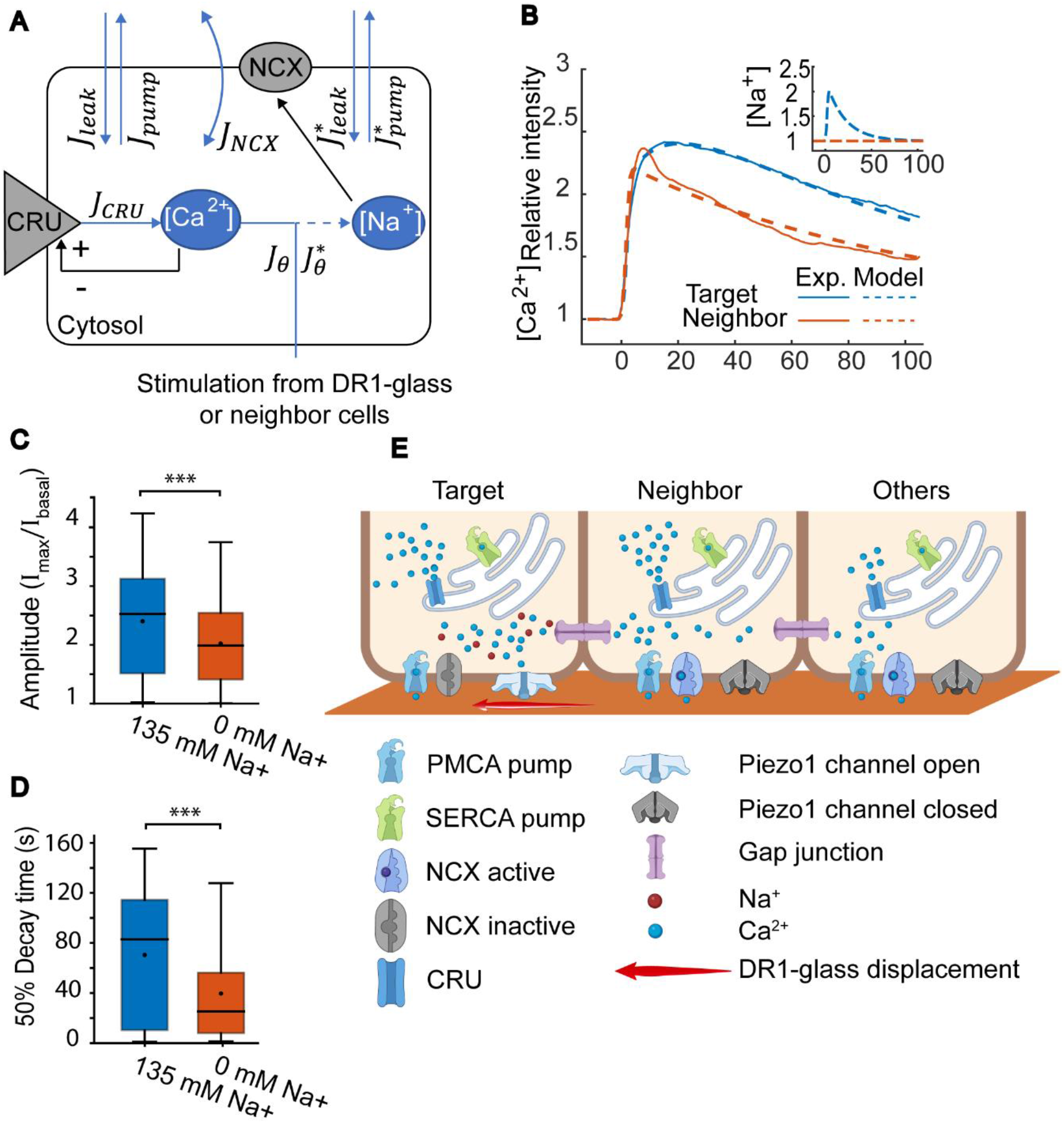
Computational model for Ca^2+^ kinetics in target cells and neighbors. **A)** Model of Ca^2+^ responses following mechanical stimulation. Blue arrows indicate the influxes and outfluxes of Ca^2+^ and N^+^ concentrations ([Ca^2+^] and [Na^+^] respectively) in cytosol. In target cells, the influx of Na^+^ is equal that of Ca^2+^ 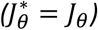. In neighbor cells, the influx of Na^+^ *J_θ_* is set to 0. **B)** The fitting of the model to the data. The averaged intensity traces of [Ca^2+^] for target cells (red) and for neighbor cells (blue) from the data (solid lines). Only the traces with amplitude of ~2.4 were selected to demonstrate that the difference in the decay time is not caused by the response amplitude alone. The normalized intensity traces predicted by the fitted models (in panel F) are shown in dashed lines in for the respective cell groups. The Na^+^ concentrations used in the model are shown in the inset panel for target cells (blue solid line) and neighbor cells (red solid line) as predicted from the fitted model. **C)** Amplitudes and **D)** decay times of cell responses in 135mM Na^+^ containing Elliot buffer and in a buffer where Na^+^ is replaced with Rb^+^. Mann-Whitney U-test indicates the statistical significance. **E)** Schematic describing the suggested signaling cascade. Shear from DR1-glass displacement is sensed in target cells by Piezo1 channels, releasing Ca^2+^ and Na^+^ into the cytosol. CRUs are activated via CICR, thus amplifying the signal from the ER. Ca^2+^ spreads to neighbor cells and other cells via gap junctions, and Ca^2+^ is amplified via CICR. [Ca^2+^] is restored by SERCA and PMCA pumps and NXCs. In target cells the NCXs are inactivated due to the influx of Na^+^. **B-C)** Boxes mark the 25% - 75% interquartile ranges (IQR), whiskers mark the range within 1.5 x IQR, horizontal lines mark the median and circles mark the mean. Statistical significances are shown as ** = p-value ≤ 0.01 and *** = p-value ≤ 0.001.

For the detailed model, please see Supporting Information section B.

From single-region stimulation experiments, we separated the Ca^2+^ traces into different groups based on quantiles of the amplitude distribution. For each group, we calculated the averaged normalized intensity (shown in Figure S5B-H, solid lines) and fit our model to these averaged traces. The only free parameters varying between the traces pertain to the influx of Ca^2+^ in cytosol (i.e., its timing, duration, and intensity). The fit results for each trace can be found in Supporting information text and Figure S5B-H. Once the effect of Na^+^ was incorporated, the model produced similar Ca^2+^ traces as the experimental data, thus supporting our hypothesis (Figure 6B).

We further tested this hypothesis of NCX involvement by performing multi-region stimulations to cells in 135 nm [Na^+^] Elliot buffer and in a similar buffer where Na^+^ was replaced with Rb^+^ (Figure 6C-D). Simply removing Na^+^ would have drastically altered the membrane potential and osmolarity of the media, leading to cell death. Therefore, it was replaced with another monovalent ion Rb^+^, that cannot be utilized by the Na^+^ transporters including NCX. When we used the Na^+^ free medium, we saw a decrease in both amplitude and especially in decay time. The mean amplitude decreased only slightly from 2.4 to 2 a.u., whereas the decay time decreased substantially from 71 s to 40 s. This demonstrates that Na^+^ affects the effectiveness of pumping Ca^2+^ out of the cytosol, and therefore supports our model.

We acknowledge that there are alternative hypotheses aside the involvement of Na^+^ to explain the different Ca^2+^ dynamics in target and neighbor cells. According to (Gottlieb, Bae, and Sachs 2012), sufficiently high stimulation can render the Piezo1 channels in a non-inactivating state, where the opening state is also a steady state instead of being transient. However, this steady state opening was only found to last at most 10s, significantly shorter than the decay time of the Ca^2+^ signals measured here. Another possible mechanism explaining the signal spreading is via Ca^2+^ induced contraction, which is a well-known phenomenon in epithelia and vascular systems (M. J. Berridge, Lipp, and Bootman 2000; Nakayama and Tanaka 1993; Takeuchi et al. 2020): the release of Ca^2+^ to cytosol due to stimulations from DR1-glass can trigger cell to contract, which exert mechanical force to neighbor cells and trigger Ca^2+^ response in them mechanically. However, we believe that the spreading time (within 2s) is too short for Ca^2+^-induced contraction (on the order of minutes (Martínez-Ara et al. 2021)) to take place. This is further supported by optogenetic activation of cell contractility, where actomyosin machinery is directly activated via light pulse. The following cell contraction happens usually in minutes, not in seconds (Cavanaugh et al. 2020).

Our data shows that also cells outside the photostimulated region exhibit Ca^2+^ transients within less than 2 seconds after the directly stimulated cells. This suggests that fast cell-cell communication occurs following the mechanical stimuli of target cells, and therefore points toward gap junction-mediated signaling. However, different signal decay kinetics was observed between the target and neighbor cells. Based on computational modeling and experimental data, we suggest that this difference is explained by the Piezo1-mediated Na^+^-influx that only occurs in target cells, and renders the NCX channels less effective, leading to slowed decay kinetics. Figure 6E portrays this distinctive effect of Na^+^ to Ca^2+^ kinetics and summarizes also all other findings presented in this research article.

## Conclusions

In this study, we demonstrated the prospects of using photoreactive azobenzene-containing materials to mechanically stimulate cells with fine spatiotemporal control. DR1-glass is often used as a convenient alternative material for one-step photolithography of micro-or nanotopographic features, recently demonstrated to be effective in controlling cell morphology and migration (C Fedele, Netti, and Cavalli 2018). However, the major benefit of using such material family is that it allows *in situ* surface manipulation in the millisecond scale with light, which is well tolerated by living cells. In this work, the material displacement upon laser scanning was thoroughly characterized in dry and wet conditions and exploited for the mechanical stimulation of epithelium. We showcased the possibility for nearly simultaneous detection of cellular responses by studying fast local and global Ca^2+^ signals in epithelial cells. Moreover, we demonstrated the key role of mechanosensitive Piezo1-channels in the sensation of basal mechanical stimulation. Particularly, we showed that instead of sensing the height of the resulting topography change, Piezo1-channels responded to the speed of shear deformation experienced by the cells, which is caused by the sideways flow of DR1-glass. The cells were able to sense even 1.48 nm/ms deformations during our 27.7 ms stimulation period. This gives new insight into the function of Piezo1-channels as well as to the dynamic interaction between epithelial cells and the ECM. Based on recent work, the basal membrane is unraveling to be an important hub for mechanical signaling in epithelial cells (Ellefsen et al. 2019; Poole et al. 2014). The dynamic interactions of the cell-ECM interface are known to guide cytoskeletal rearrangements such as cell attachment, orientation, and migration, as well as gene expression (Martino et al. 2018). In larger scale these interactions are related to epithelial signaling during development and differentiation (Sui et al. 2018). However, the molecular effects of these mechanically induced Ca^2+^ transients are yet unknown. Interestingly, the localization of Piezo1 channels (on the basal membrane) consequently also place the Ca^2+^ signals there in proximity of the actin cytoskeleton. This suggests a possibility for a Ca^2+^-based feedback loop between the dynamic ECM and the actively remodeling cytoskeleton. More importantly, when malfunctioning, dynamics in this interface have a significant role in cancer progression (Reuten et al. 2021; Niland, Riscanevo, and Eble 2021). As over 80 % of cancers originate from epithelial tissue, the importance of understanding these cell-ECM interactions is evident. Interestingly, Piezo1 channels are known to be overexpressed in several cancers (De Felice and Alaimo 2020), thus our work gives insight and provides a promising tool for studying epithelial cancer progression.

In addition, we demonstrated the spatial flexibility of the system that enables a mechanical stimulation ranging from local subcellular probing to population wide mechanical activation. We used this flexibility to study mechanically induced intercellular Ca^2+^ signaling. We identified two types of signal kinetics distinguishing between the mechanically targeted cells and the neighbor cells where the signal spreads. We developed an experimentally verified computational model that explains the differing kinetics by distinctive involvement of Na^+^. Lastly, we showed that within a cell population there is a balance between cell-to-cell communication and direct mechanical activation, leading to heterogeneity in the responses. Our findings are summarized in Figure 6E.

So far, the focus of our work has been on the material deformation triggered Ca^2+^ signaling, with the signals’ time scale on the order of ~100s. These Ca^2+^ transients can invoke several longer-term changes in cells, e.g., contraction, proliferation, migration. It should be noted that cells may also concurrently adapt to changed physical environment via these processes, e.g., DR1-glass photoinscribed topography (Fedele et al. 2020). Cellular adaptation needs to be taken into consideration when studying the long-term impacts of the mechanical stimuli. However, the possible causality between these two mechanical signaling processes, the fast Ca^2+^ signals and long-term impacts such as orientation and migration, remain unknown. Our future work aims to focus on this possible relationship.

## Supporting information

Support information

## Acknowledgments

This research was supported by Emil Aaltonen project funding, Academy of Finland Research Fellow funding (308315, 314106), Academy of Finland Post-Doctoral funding (332615), Academy of Finland consortium project BioMiRI (323507), LIBER Center of Excellence (346107), Academy of Finland PREIN Flagship program (320165), and Tampere University Health Data Science profiling area’s Post-Doctoral fellowship (2021). The authors acknowledge the Biocenter Finland (BF), Tampere Imaging Facility (TIF) and the Tampere facility of Flow Cytometry for their service. Douglas Kim & GENIE Project are gratefully acknowledged for the pGP-CMV-NES-jRCaMP1b plasmid. Mari Isomäki is acknowledged for her help. Figures 3A and 5J were created with BioRender.com.

## Materials and Methods

### DR1-glass sample preparation

Glass coverslips (22×22 mm^2^) were washed in acetone in a sonicating bath and spin coated with a solution of 3,25 % (g·ml^-1^) of Disperse Red1 molecular glass (DR1-glass, Solaris Chem. inc., Canada) in CHCl3. The spin coating parameters were 1500 rpm for 30 seconds.

### Cell culture

A stable MDCK II cell line expressing the red calcium indicator jRCaMP1b (Dana et al. 2016) was established. The Neon electroporation system was used to transfect cells. Positive colonies were first enriched with 600 μg/ml G418 (4727894001, Sigma-Aldrich) selection and finally FACS sorting was used to generate a positive cell line. The plasmid pGP-CMV-NES-jRCaMP1b was a gift from Douglas Kim & GENIE Project (Addgene plasmid #63136; http://n2t.net/addgene:63136; RRID: Addgene_63136). MDCK II jRCaMP1b cells were maintained in Minimum Essential Medium (MEM) with GlutaMAX™ and Earle’s salts (41090-028, Gibco), supplemented with 10 % FBS (10500064, Gibco) and 250 μg/ml G418 (#4727894001, Roche, Switzerland). Penicillin streptomycin was not used as it is a competitive inhibitor for G418. Media was changed twice a week and cells were passaged every 7-14 days.

For cell experiments DR1-glass samples were coated with fibronectin. Fibronectin (purified from human plasma) was diluted to 10 μg/ml in Dulbecco’s Phosphate Buffered Saline (PBS) and the coating solution was pipetted on top of samples at approximately 150 μl/cm^2^. Samples were air dried under UV light for 45 min and washed twice with PBS before adding media and cells. Cells were plated at approximately 3.5 x10^4^ cells/cm^2^ and cultured for 6 days to reach mature epithelium at optimal cell density. Cells were allowed to stabilize in the microscope chamber with +37°C incubation and 5% CO_2_ gas flow for approximately 30 minutes before commencing imaging.

As negative control sample, we used a DR1-glass sample flipped upside down (Figure 3A). Therefore, the absorbance spectrum was identical to DR1-glass samples, but cells were not in contact with the photopatterning. Fibronectin coating was performed as previously explained, and cells were cultured for the same 6-day period. However, 1.75 x 10^4^ cells were plated to reach comparable cell density.

### Photopatterning and detection of calcium signals

Material photopatterning was carried out at LSM 780 laser scanning confocal microscope (Zeiss Microscopy, Germany) equipped with a large incubator with heating and CO_2_-control. The samples were stimulated with a water immersion objective, Zeiss C Apo 63x/1.20, WD 0.28 mm with a 488 nm Argon laser in photobleaching mode. Briefly, transmitted light (T-PMT) was used to focus the stimulation laser to the DR1-glass layer and defined regions of interest (3 x 5 rectangles á 7×400 pixels) were drawn over the surface with unidirectional scanning mode. A 512×512 field of view was imaged with 200 nm pixel size. Pixel dwell time was 1.00 μs and frame rate for imaging 1.23 s. The pattern inscription time per one rectangle was 27.7 ms. Samples were placed into AirekaCells^®^ coverslip cell chambers (SC15022, Aireka Scientific Co., Ltd, Hong Kong) and either imaged dry, with 1ml deionized water, or with 1 ml of conditioned media on top.

For material characterization cell free DR1-glass samples were used. Photopatterning was performed both to dry samples and to fibronectin coated samples with media on top. 4.8, 7.4, 9.5, 11.7, and 14.4 mW/cm^2^ laser intensities with 1, 5, 10, 20 and 30 iterations were tested. Each combination of parameters was repeated 3 times. The laser intensities were measured at the objective back focal plane are reported in Figure S1 in Supporting Information.

For cell experiments, we chose areas with normal cell morphologies and jRCaMP1b expression, and where no fluid filled dome structures were apparent. 561 nm excitation was used to detect calcium and T-PMT to focus the photobleaching laser to the DR1-glass layer. Photopatterning was produced with the same 488 nm laser as described before. To ensure optimal imaging of the calcium indicator and stimulation of the DR1-glass, different focal levels were used for 561 nm and 488 nm channels. The spontaneous baseline activity of cells was recorded for 10 frames after which stimulation was performed as described above. Following the stimulation, the cell responses were recorded for 121 frames. 4.8, 7.4, and 9.5 mW/cm^2^ laser intensities with 1, 5 and 10 iterations were used. In addition to the multi-region stimulation described above, also single-region stimulation (a single 7×400 pixel rectangle) was performed.

### Pharmaceutical experiments

For pharmaceutical experiments cells were cultured and samples prepared as described previously. Cells were first imaged in normal conditions as described before to acquire control data. Thapsigargin (T9033, Sigma-Aldrich), Yoda1 (5586, Tocris Bioscience, Bristol, UK), and cytochalasin D (C2618, Sigma-Aldrich) were dissolved in DMSO and GsMTx4 (4912, Tocris Biosciences) in H_2_O. Thapsigargin was used at 1 μM, Yoda1 at 10 μM, cytochalasin D at 10 μg/ml, and GsMTx4 at 10 μM. Drugs were added on the sample and gently mixed. After addition of the pharmaceutical, cells were allowed to relax for 15 min – 1 h before repeating stimulation experiments in the presence of the pharmaceutical.

### Characterization of surface topographies

The samples were characterized with UV/Vis spectrophotometer (Cary 60 UV/Vis spectrophotometer, Agilent, California). The surface topographies were characterized with atomic force microscopy (Park XE-100 AFM, Park Systems, Korea) in non-contact mode in air with an Al-coated Si ACTA probe (Applied NanoStructures, California) with 200–400 kHz nominal frequency and 13–77 N m^−1^ spring constant. Digital Holographic Microscopy (DHM R-2100, Lyncée tech., Switzerland) was used to image large areas of the sample and obtain quantitative information about the photoinscribed surface reliefs.

### Analysis of material displacement

To follow the material displacement, fluorescent microbeads were deposited over the material surface (1% solid content, FluoSpheres™ Carboxylate-Modified Microspheres, 0.2 μm diameter, dark red fluorescent (660/680), Invitrogen, ThermoFisher Scientific, Massachusetts) for 1 h. The microspheres were imaged with a 633 nm diode laser in a time-lapse alternating bleaching and imaging frames. For the analysis of material light-induced displacement in absence of a water-based medium, Particle Image Velocimetry was used. The analysis was carried out using PIVlab MATLAB plugin on consecutive image pairs from the transmitted light channel (Thielicke and Sonntag 2021). The analysis was performed using a 3-pass analysis with first pass interrogation window of 64 × 64 pixels with a 50% overlap. The material flow was extracted as frame average value within each region of interest from smoothed vector fields via a custom MATLAB script. The light-induced displacement of the material in presence of water was quantified by measuring in Fiji ImageJ software (Schindelin et al. 2012) the distance covered by the fluorescent particles during the photopatterning experiment.

### Calcium signal analysis

Calcium signals were analyzed with Fiji ImageJ software and Matlab. Regions of interest (ROI) were segmented with Fiji ImageJ by manually fitting ellipses around the nuclei and by expanding the ellipses into 1 μm thick bands surrounding the nuclei. A custom Matlab script is used to extract the calcium cytosolic concentration in each cell as the mean intensity value of each ROI in each frame of the time lapse. This time traces of calcium signals are then normalized by the baseline intensity (I_basal_, calculated from the first 10 frames prior to photostimulation). From each trace, we calculated the Amplitude as the maximum of the normalized signal intensity (I_max_/I_basal_), and the decay time as the time from the maximum to mid-level between the maximum and basal levels. Independent samples Mann-Whitney U test was used for statistics when comparing the distributions of Amplitudes and decay times between conditions.

### Immunostainings, imaging, and image processing

Cells were fixed with 1% (for Piezo1) or 4 % (for phalloidin) paraformaldehyde (PFA, Electron Microscopy Sciences 15713-S) for 10 minutes and washed two times with PBS. Immunostainings were performed in RT protected from light. Samples were permeabilized with permeabilization buffer (0.5% BSA and 0.5 % Triton-X 100 in PBS) for 10 min before blocking with 3 % BSA (PAN-Biotech P06-139210) in PBS for 1h. The primary antibody (Piezo1 NBP1-78446) was diluted 1:50 in blocking buffer and incubated for 1h. Samples were washed 3 x 10 min (permeabilization buffer, 1x PBS, permeabilization buffer) before adding the secondary antibody (Anti-Anti-rabbit-488 A11008 and phalloidin-647, or only phalloidin-488) diluted (anti-rabbit 1:200 and phalloidins 1:100) in blocking buffer for 1h. Samples were washed 2 x 10 min with PBS and 5 min with deionized H_2_O before mounting with Prolong Diamond with DAPI (P36962, Thermo Fisher Scientific, Waltham, MA USA). The immunostained samples were stored in +4°C protected from light.

Immunostained samples were imaged with laser scanning confocal microscope Nikon A1R mounted in inverted Nikon Ti-E body (Nikon, Tokyo, Japan), with SR Apo TIRF 100x/1.49 objective. Pixel size was set to 40 x 40 nm and Z-step to 99 nm. Deconvolution was performed with Huygens Essential (Scientific Volume Imaging, Hilversum, Netherlands) with automated standard settings. FIJI was used to make maximum intensity projections of z-stacks and to adjust image brightness and contrast.

### Na^+^-free media tests

The effect of Na^+^ to decay time of calcium signals was tested with Elliot buffer either containing Na^+^ or with Na^+^ replaced by Rb^+^. The full formulation was 137 mM NaCl/RbCl, 5 mM KCl, 1.2 mM MgCl_2_, 2 mM CaCl_2_, 0.44 mM KH_2_PO_4_, 4.2 mM NaHCO_3_, 20 mM HEPES, 5 mM glucose, and 5% FBS. The pH was adjusted to 7.4 and the osmolarities were 317 mOsm/l (Elliot + Na^+^) and 301 mOsm/l (Elliot + Rb^+^). Samples were coated with fibronectin and cells cultured in MEM as described before. Before imaging, the media was first changed to Elliot + Na^+^ and 3-4 stimulation experiments were performed as described before. Then media was changed to Th Elliot + Rb^+^ and another 3-4 stimulation experiments were performed. Independent samples Mann-Whitney U test was used for statistical testing.

### Apical micromanipulation

Imaging was done with Nikon Eclipse FN1 widefield fluorescence microscope (Nikon, Tokyo, Japan) equipped with Andor DU-888 X-11486 camera with NIR Apo 40x 0.8W DIC N2 objective. 558 nm wavelength excitation light was used. Sensapex (Oulu, Finland) uMp-RW3 micromanipulator was used to control the patch pipette (resistance 4-8 MΩ) containing Ames’ solution (A1420, Sigma Aldrich) buffered with 10mM HEPES. A target cell was chosen, and during imaging the cell was approached until a calcium response was seen. 2 min videos were acquired.

### Model of Calcium dynamics

The model of calcium dynamics is extended from the simplified model in Kaouri et al. (Kaouri et al. 2019) to account for the role of Na^+^ cytosolic concentration and Na^+^/Ca^2+^ exchanger (NCX) (Figure 6A). The model is deterministic, with only the amplitude of the calcium influx into cells’ cytosol following mechanical stimulations *J_θ_* allowed to vary between cells. Before model fitting, the calcium traces in each cell type (target or neighbor) are split into different subgroups based on quantiles of the signals’ amplitude. The model parameters are then learned from the averaged calcium signal intensity in each subgroup. Please see Supporting Information section B for the detailed model assumption and the fitting process.

## Notes

### Competing Interest Statement

The authors have declared no competing interest.

### Summary of Updates

Updated with small editing on acknowledgements and links to raw data

