## Supplementary material for "Light-induced nanoscale deformation in azobenzene thin film triggers rapid intracellular Ca^2+^ increase via mechanosensitive cation channels": Support information

### Supporting information text

**A. Determine the effect of amplitude on 50% decay time in calcium responses**

We create a mathematical framework to assess whether the difference in the 50% decay time of the calcium responses between two cell populations is caused by changes in the responses' amplitude alone. From two cell-populations, we obtain two set of samples $(x_{1i}, y_{1i})_{i=1..M}$ and $\left(x_{2j}, y_{2j}\right)_{j=1..N}$, each sample contain a pair of values corresponding to the calcium response' 50% decay time $(x)$ and amplitude $(y)$ in each cell. The amplitude distributions in cell population 1 and 2 are given by $P(Y_1)$ and $p(Y_2)$ respective.

The conditional distribution of $X_1$ the 50% decay time given each value of the amplitude in population 1 $Y_1$ is given by:

$$p(X_1 = x|Y_1 = y) = p(X_1 = x, Y_1 = y)/p(Y_1 = y) \quad (1)$$

Here, we compute $X_1'$ the distribution of the 50% decay time in the population 1, if their amplitude follows the distribution in population 2.

$$p(X_1' = x) = \int_{y=0}^{\infty} \frac{p(X_1 = x, Y_1 = y)p(Y_2 = y)}{p(Y_1 = y)} dy \quad (2)$$

For simplicity, we bin the values of amplitude $y_{1i}$ and $y_{2j}$ and values of 50% decay time $x_{ij}$ and $x_{2j}$ to the nearest values in the $\epsilon_Y$ and $\epsilon_X$, respectively. $\epsilon_Y$ contains 10 equally-spaced bin values from 0 to the maximum of amplitude 2.6. $\epsilon_X$ contains 10 equally-spaced bin values from 0 to the maximum of 50% decay time 165 s. Therefore, the probability density functions $p()$ can be approximated by the mass density functions $P()$. The probability mass function of $X_1'$ is given by:

$$P(X_1' = x) = \sum_{y \text{ in } \epsilon_Y} \frac{P(X_1 = x, Y_1 = y)P(Y_2 = y)}{P(Y_1 = y)} \quad (3)$$

Here, $P(X_1 = x, Y_1 = y)$, $P(Y_1 = y)$ and $P(Y_2 = y)$ can be calculated directly from the binned data. Therefore, the approximated probability of 50% decay time in population 1 $X_1'$ if the amplitude follows $Y_2$'s distribution can be calculated from Eq. 3 and compared directly with $P(X_2)$.

In our work, $X_1'$ is the expected 50% decay time from calcium responses in Neighbor cells if the amplitude distribution follows that in Target cells (Figure 5D). This allows us to directly compare the kinetics of signal decay in Target $(X_2)$ and Neighbor cells $(X_1')$, independent of the response amplitude (Figure S6A).

**B. Model description and parameter estimation**

#### Model of Ca2+ response in directly and non-directly stimulated cells

We extend the simplified model of calcium ($Ca^{2+}$) response in Kaouri et al. (Kaouri et al. 2019) to account for $Na^+$ cytosolic concentration (*Fig. 5D*). In this model, the dynamics of $Ca^{2+}$ cytosolic concentration $c$ is defined as:

$$\frac{dc}{dt} = J_\theta(t) + J_{CRU} + J_{pump} + J_{leak} + J_{NCX}. \quad (4)$$

In Eq. 4, $J_\theta$ is the influx of $Ca^{2+}$ from the extracellular environment (either via gap junctions or from the media). $J_{CRU}$ is the calcium influx from Calcium Release Unit (CRU). $J_{pump}$, $J_{NCX}$ are the calcium influx induced via SERCA pumps and $Na^+/Ca^{2+}$ (NCX) exchanger respectively. $J_{leak}$ is the leaky flux from either the extracellular environment or intracellular calcium storages.

We assume that following the stimulation applied at time $t_0$ (in seconds), $J_\theta(t)$ depends on the maximum $Ca^{2+}$ passage rate through stimulated channels $k_\theta$ (either directly through piezo channels or indirectly via $Ca^{2+}$ specific channels) and the average time these channels are opened $\tau_\theta$ . To ease the stiffness in ODE simulation (occurring when sudden fluxes are arbitrarily introduced), we set the fraction of opened channels over time following a gaussian curve, peaked at $t_0 + 2\tau_\theta$:

$$J_\theta(t) = k_\theta \frac{1}{\sqrt{2\pi}} e^{-\frac{(t-t_0-2\tau_\theta)^2}{2\tau_\theta}} \quad . \tag{5}$$

From Eq. 5, the total $Ca^{2+}$ flux from the external environment is:

$$\int J_\theta(t)\, dt = k_\theta \tau_\theta. \tag{6}$$

The Calcium Release Unit (CRU) can be both activated and inactivated by $Ca^{2+}$ at different concentration.

$$J_{CRU} = k_{CRU} f_1 f_2, \tag{7}$$

where $f_1$ and $f_2$ are respectively the fraction of Ca2+ activated and inactivated CRU:

$$f_1 = \frac{c^{h_1}}{c^{h_1} + K_1^{h_1}}, \tag{8}$$

$$f_2 = \frac{K_2^{h_2}}{c^{h_2} + K_2^{h_2}}. \tag{9}$$

In Eq. 8 and 9, $K_1$ and $K_2$ are respectively the 50% activating and inactivating thresholds for $Ca^{2+}$ cytosolic concentration $c$. $h_1$ and $h_2$ are respectively the order of the Hill functions describing the CRU's activation and inactivation. For the system to be both monostable and with hysteresis, we set $h_2 = 4$ and $h_1 = 3$.

$J_{pump}$ is the total outflux of $Ca^{2+}$ being pumped out of the cytosol (e.g., via SERCA pumps or PMCA pumps (Blaustein and Lederer 1999; Jensen, Buckby, and Empson 2004)), independent of cytosolic $Na^+$ concentration:

$$J_{pump} = -k_{pump} c. \tag{10}$$

$J_{leak}$ is the leakage of $Ca^{2+}$ from the external environment:

$$J_{leak} = \beta. \tag{11}$$

$Ca^{2+}$ is also pumped in and out of cells via $Na^+/Ca^{2+}$ exchanger (NCX), which depends on the cytosolic $Na^+$ concentration $n$:

$$J_{NCX} = k_{NCX+} \frac{n^3}{n^3 + K_n^3} - k_{NCX-} c, \tag{12}$$

with $k_{NCX+}$ and $k_{NCX-}$ are the respective import and export rate constants of $Ca^{2+}$ into and out of cytosol via NCX. We assume that the external concentration of $Na^+$ and Ca2+ is abundant and constant throughout the experiments and thus omitted in the value of $k_{NCX+}$ and $k_{NCX-}$. The order of the $Ca^{2+}$ import rates as a function of $n$ is set to 3 to account for the three $Na^+$ ions required to import a single $Ca^{2+}$ ions (Matsuoka and Hilgemann 1992). Note that in Eq. 10 and 12, it is impossible to distinguish between the effects of changing $k_{pump}$ and $k_{NCX-}$. Therefore, for convenience, we assume $k_{NCX-} = 0$:

$$J_{NCX} = k_{NCX+} \frac{n^3}{n^3 + K_n^3}. \tag{13}$$

The dynamics of $Na^+$ concentration $n$ is given by:

$$\frac{dn}{dt} = J_\theta^*(t) + J_{pump}^* + J_{leak}^*, \tag{14}$$

in which $J_\theta^*(t)$ is the influx $Na^+$ from the extracellular environment. For directly stimulated cells, as piezo channels are opened allowing passive diffusion of ions across cell membranes, $J_\theta^*(t)$ depends on the average open duration of piezo channels following stimulation $\tau_\theta$ (as in Eq. 3). As diffusion of ions through piezo channels are passive, for convenience, we assume that on the same maximum passage rate through piezo channels for $Na^+$ and $Ca^{2+}$:

$$J_\theta^*(t) = J_\theta \text{ for directly stimulated cells.} \tag{15}$$

For indirectly stimulated cells, we assume that there is no influx of $Na^+$. Therefore:

$$J_\theta^*(t) = 0 \text{ for indirectly stimulated cells.} \tag{16}$$

$J_{pump}^*$ is the total outflux of $Na^+$ being pumped out of the cytosol via e.g., $K^+/Na^+$ exchangers, given by:

$$J_{pump}^* = -k_{pump}^* n. \tag{17}$$

Note that in Eq. 17, we don't take into account outflux of $Na^+$ via NCX (Eq.9). As the $Na^+$ cytosolic concentration (on the order of µM (Willumsen and Boucher 1991)) is much higher than that of $Ca^{2+}$ (on the order of nM) in epithelial cells in resting conditions, exchanges via NCX should only affect the latter's level.

$J_{leak}^*$ is the leakage of $Na^+$ from the external environment:

$$J^*{}_{leak} = \beta^*. \tag{18}$$

#### Parameter calibration

In our experiments, $Ca^{2+}$ levels are measured only with a fluorescent reporter and do not inform about the absolute concentration. Therefore, we focus on explaining the relative changes in $Ca^{2+}$ concentration following perturbations, i.e., the normalized $Ca^{2+}$ and $Na^+$ levels is unity ($c = 1, n = 1$) and at steady state, prior to any mechanical stimulations ($t = 0$):

$$\begin{aligned}
&\frac{dc}{dt} = J_{CRU} + J_{pump} + J_{leak} + J_{NCX} = 0 \mid c = 1, n = 1 \\
&\leftrightarrow k_{CRU} f_1 f_2 - k_{pump} c + \beta + k_{NCX+} \frac{n^3}{n^3 + K_n^3} = 0 \mid c = 1, n = 1 \\
&\leftrightarrow k_{CRU} \frac{1}{1+K_1^{h_1}} \frac{K_2^{h2}}{1+K_2^{h2}} - k_{pump} + \beta + k_{NCX+} \frac{1}{1+K_n^3} = 0 \\
&\leftrightarrow k_{CRU} \frac{1}{1+K_1^3} \frac{K_2^4}{1+K_2^4} + k_{NCX+} \frac{1}{1+K_n^3} + \beta = k_{pump}.
\end{aligned} \tag{19}$$

$$\begin{aligned}
&\frac{dn}{dt} = J_{pump}^* + J_{leak}^* = 0 \mid c = 1, n = 1 \\
&\leftrightarrow k_{pump}^* = \beta^* \ .
\end{aligned} \tag{20}$$

With the constraints from Eq. 19 and Eq. 20, we can reduce the model's 9 parameters to 7 free parameters to model the resting condition. To model the influx of ions following stimulation (either direct or indirect), we use three parameters $t_0$, $k_\theta$ and $\tau_\theta$.

#### Fitting the model to the data

To quantify the effect of $Na^+$ influx on the $Ca^{2+}$ response, we fit the model to $Ca^{2+}$ traces measured with jRCaMP1b in direct and indirectly stimulated cells. As $Ca^{2+}$ responses vary highly between cells in term of amplitude, we first separate the traces by 5-quantiles based on the normalized amplitude distribution into groups (5 groups from Target cells and 5 from Neighbor cells). From each $i^{th}$ of the 10 groups, the averaged normalized intensity $I_i(t)$ at time $t$ is computed (Fig. S6). Given the microscopy's total duration (120 s) and frame rate (1 frame/s), $t$ takes integer values from 1 to 120 (unit in second).

When fitting the model to these averaged traces, we assume that all traces share the same parameters' values except for the average time the piezo channel is open $\tau_\theta$, the value of which can vary between trace groups and is thus the driving source of cell-to-cell variability. As the averaged amplitude from 3 weakest groups from Neighbor cells are very low (<1.1), we exclude them from the fit to reduce the number of free parameters. Therefore, only 7 trace groups $\epsilon_i = [1,2,3,4,5,9,10]$ are used for the fitting. The parameter set $X$ to be optimized is:

$$X = [\tau_{\theta,i}, k_\theta, k_{CRU}, K_1, K_2, k_{pump}, \beta, k_{NCX+}, K_n] \text{ with } i \text{ in } \epsilon_i \quad (21)$$

Given each parameter set $X$, we find the expected $Ca^{2+}$ response for the i$^{th}$ trace $I_{i,model}(t)|X$ by solving the set of ODE in Eq. 4, 9 and 14. The objective function $L(X)$ to minimize is the total squared distance between the measured and the predicted response:

$$L(X) = \sum_{i \text{ in } \epsilon_i} \sum_{t=1..120} \left(I_i(t) - I_{i,model}(t)\right)^2 \quad (22)$$

We use zeroth-order Nelder-Mead simplex direct search (Lagarias et al. 1998) for parameter estimations. To ensure that the estimated parameter set corresponds to the global minimum, we perform the optimization with 200 randomized initial values of $X$ and the optimized value of $X$ yielding the best fit is selected.

See Table 1 for a summary of the parameter description and their fitted values. The predicted curves in comparison with the data are shown in Fig. S6 for each group, along with the fitted parameter values for $k_{\theta,i}$, $\tau_{\theta,i}$.

Table 1: Models' best fitted parameters to the data. The * sign in the *fitted values* column indicates that the values are fixed and thus not optimized in the fit.

| Parameters | Description | Fitted values |
|---|---|---|
| Initial ion fluxes | | |
| $k_\theta$ | Maximum passage rate of $Ca^{2+}$ through stimulated (either via piezo or $Ca^{2+}$ specific) channels | Varying between traces |
| $\tau_\theta$ | Average duration of open channels following stimulation | Varying between traces |
| $t_0$ | Time of stimulation | 10 s* |
| Calcium Release Unit (CRU) | | |
| $k_{CRU}$ | Maximum $Ca^{2+}$ release rate from fully activated CRU | 0.079 |
| $K_1$ | Threshold for 50% CRU activation by $Ca^{2+}$ | 1.51 |
| $h_1$ | Hill coefficient for CRU activation by $Ca^{2+}$ | 3* |
| $K_2$ | Threshold for 50% CRU inactivation by $Ca^{2+}$ | 66 |
| $h_2$ | Hill coefficient for CRU inactivation by $Ca^{2+}$ | 4* |
| Ion pumps and leakages | | |
| $k_{pump}$ | maximum $Ca^{2+}$ pumping rate from cytosol | 2.81 /s |
| $k^*_{pump}$ | maximum $Na^+$ pumping rate from cytosol | 0.023 /s |
| $\beta$ | $Ca^{2+}$ leakage rate to cytosol | 0.024 /s |
| $\beta^*$ | $Na^+$ leakage rate to cytosol (equal $k^*_{pump}$) | 0.023 /s |
| $Na^+/Ca^{2+}$ exchanger (NCX) | | |
| $k_{NCX+}$ | Maximum rate of $Ca^{2+}$ import via NCX | 0.25 /s |
| $k_{NCX-}$ | Maximum rate of $Ca^{2+}$ export via NCX | 0* |
| $K_n$ | Threshold for 50% NCX activation by $Na^+$ | 1.98 |

#### Supporting information figures:

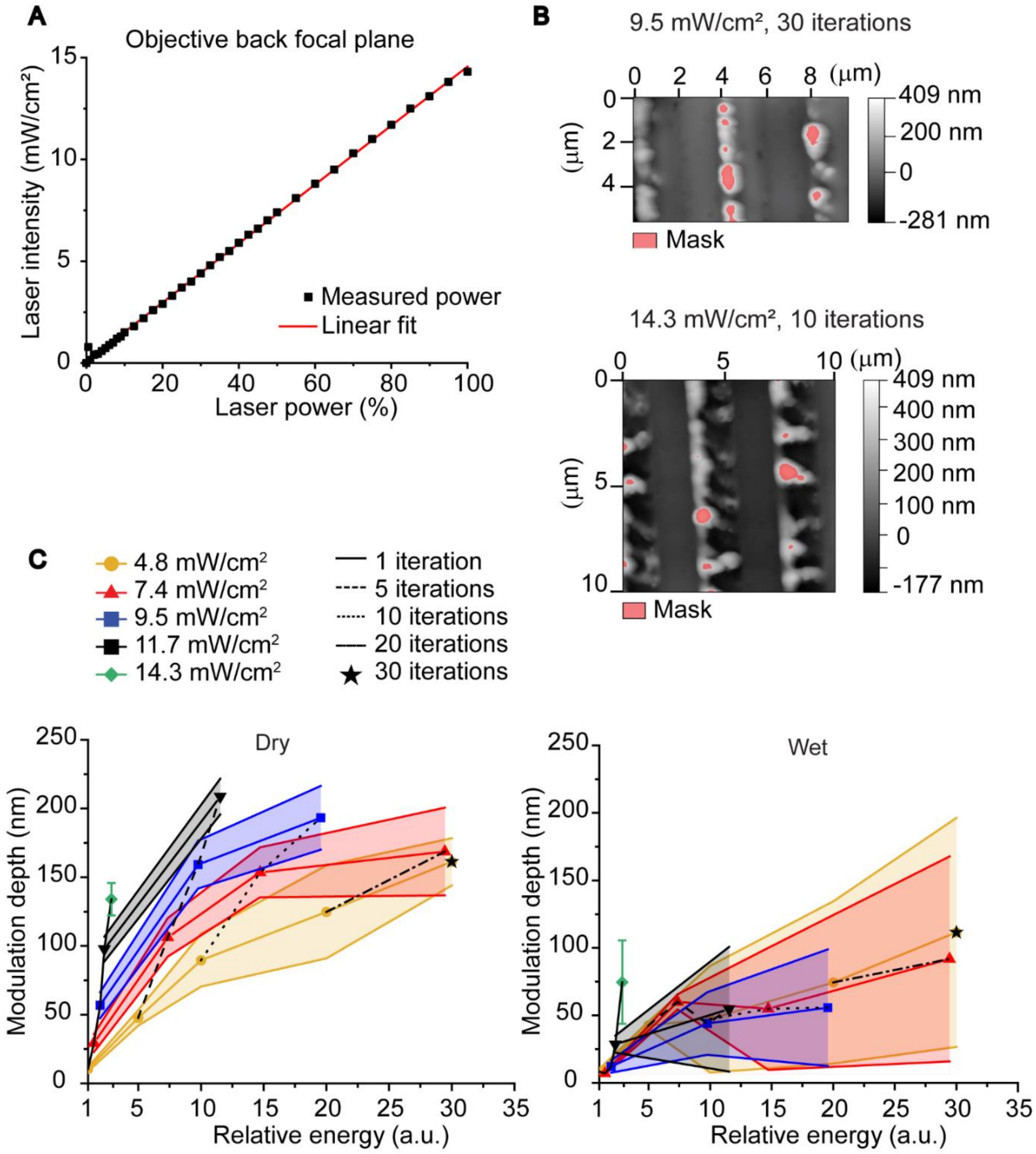

**Figure S1: Laser intensity effects on DR1-glass topography.**
*A) Laser power as measured at the back focal plane. B) AFM micrographs of light-induced topographic features with higher laser power and number of iterations. C) Modulation depth of the microtopographies and a function of the total energy dose in dry and wet conditions in the full power range used (33%-100% of the total laser power).*

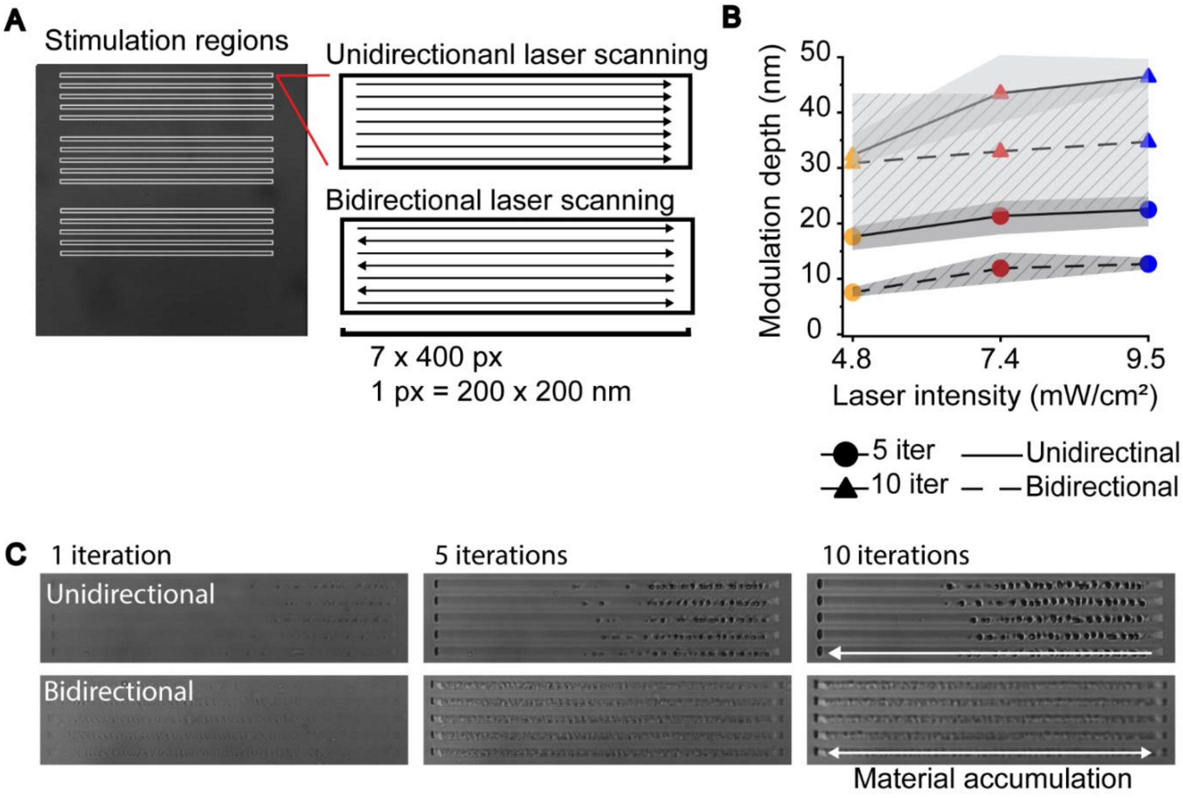

**Figure S2: Effects of scanning mode to material topography and piling.**

*A) Representation of the stimulation area and the two used scanning modes: unidirectional and bidirectional. B) Comparison of means ± StDevs of modulation depths produced with unidirectional (continuous line) and bidirectional (dashed line) scanning with 4.8 mW/cm$^2$ (yellow), 7.4 mW/cm$^2$ (red) and 9.5 mW/cm$^2$ (blue) laser intensities in wet conditions. Only data for 5 iterations (circles) and 10 iterations (triangles) are presented as the modulation depths are so low for 1 iteration. C) The piling of material after 1, 5 and 10 iterations with unidirectional and bidirectional laser scanning in 5 separate stimulated rectangles. With unidirectional scanning the material piles to the left side whereas with unidirectional scanning the piling is symmetrical at both sides.*

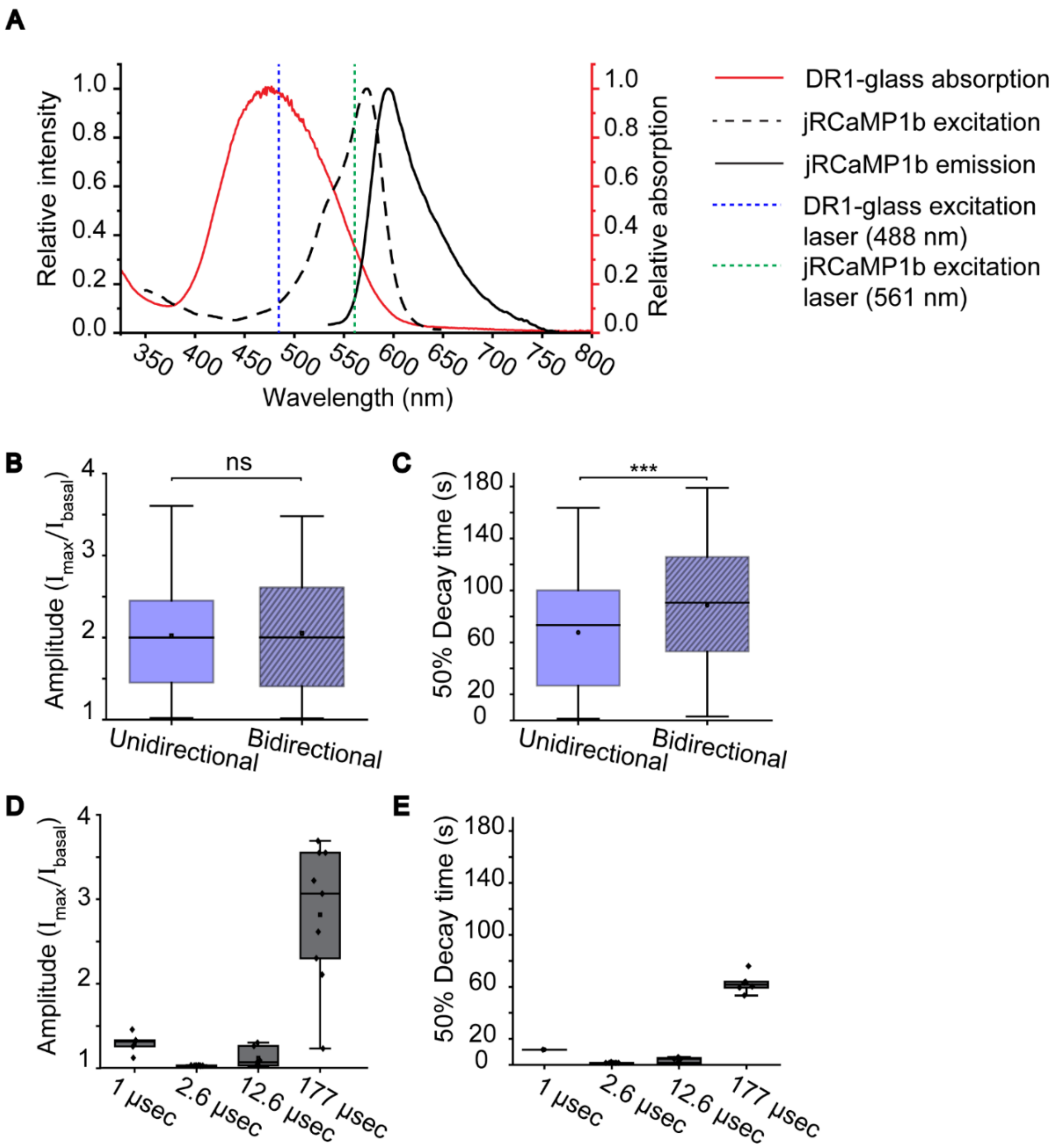

**Figure S3. Effects of scanning mode and pixel dwell time on cell responses.**

*A) The absorption spectrum of DR1-glass (red line) and the excitation (black dashed line) and emission (black solid line) spectra of the calcium indicator jRCaMP1b, and the excitation laser wavelengths for DR1-glass (488 nm, blue dashed line) and jRRCamP1b (561 nm, green dashed line) B) The amplitude and C) 50% decay time of cells stimulated with unidirectional or bidirectional (striped) laser scanning. D) Amplitude and E) 50% decay time of cells stimulates with different pixel dwell times. 9.5 mW/cm$^2$ laser intensity and 1 iteration were used in all experiments.*

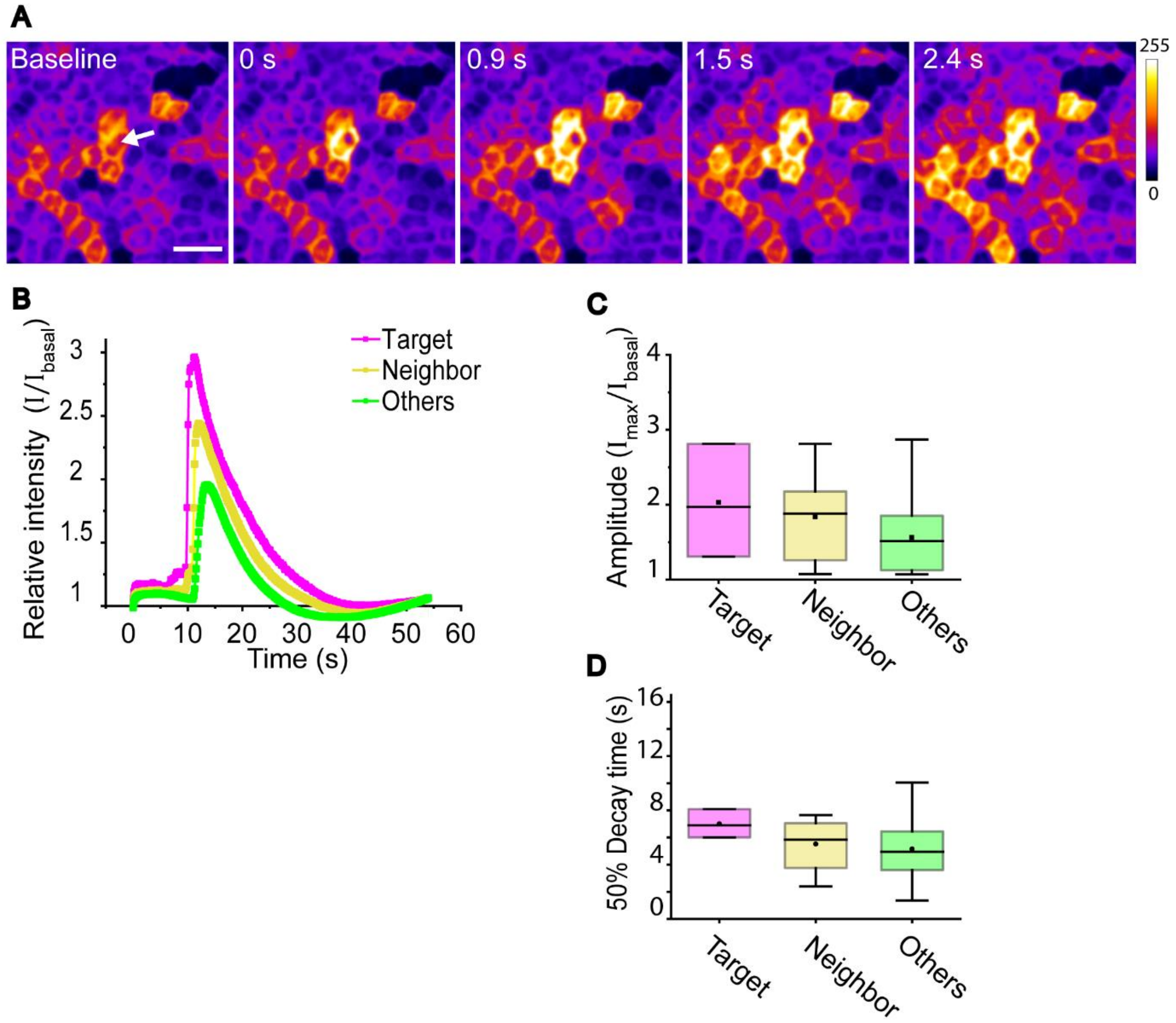

**Figure S4. Apical stimulation.**

*A) Single frames from before stimulation, immediately after stimulation and 0.9 s, 1.5 s and, 2.4 s after stimulation. The target cell is marked with a white arrow, scale bar 50 µm. Color bar represents calcium signal intensity. B) Mean ±SE intensity plots of cell responses in the different cell groups (targeted cells, neighbors, and others). C) Normalized amplitudes and D) 50% decay times of cell responses. Note that in (D) the scale is 10-fold smaller than in other similar plots.*

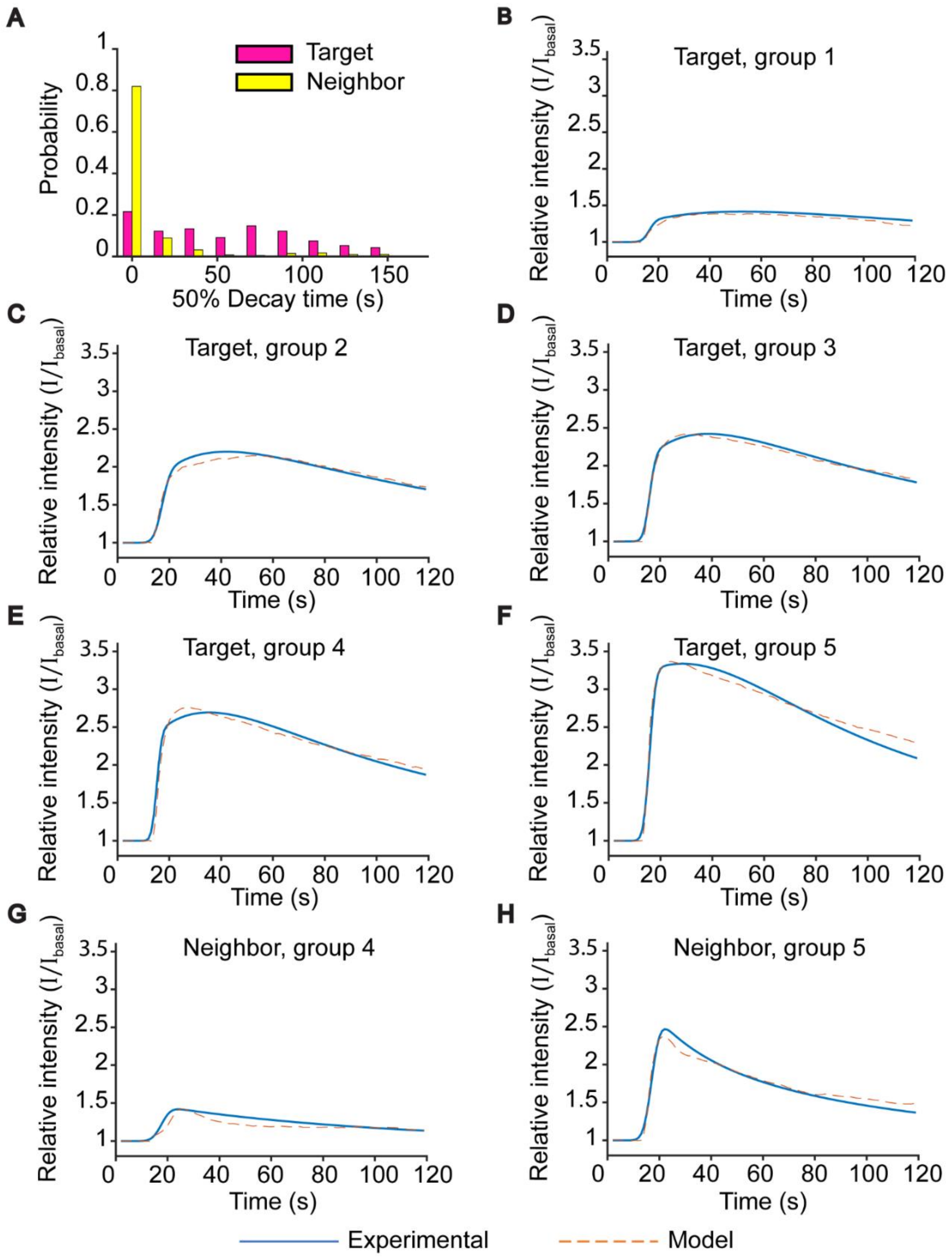

**Figure S5. Distinguishing calcium dynamics in Target and Neighbor cells:**
*A) Calibrated distribution of 50 % Decay* time in Target cells (purple bar) and Neighbor cells (yellow bar), given the same distribution of response Amplitude as in Target cells (see SI section A). **B-H)** Comparison of the mean calcium intensity in each 5-quantile groups from the experiment (solid blue curves) vs fitted model's prediction (dashed orange curves). The response's amplitude in group 1 to 3 in Neighbor cells is very small (<1.1) and thus not used for the fit.

#### Supporting information videos:

**SI Video 1: Unidirectional scanning**
**SI Video 2: Bidirectional scanning**
**SI Video 3: Calcium responses in multiline stimulated sample in normal conditions**
**SI Video 4: Calcium responses in multiline stimulated sample with Yoda1 (10 µM)**
**SI Video 5: Calcium responses in multiline stimulated sample with Cytochalasin D (10 µg/ml)**
**SI Video 6: Calcium responses in multiline stimulated sample with Thapsigargin (1 µM)**
**SI Video 7: Apical stimulation**

**All videos can be accessed via:** https://doi.org/10.5281/zenodo.7138621
